# Force-transducing molecular ensembles at growing microtubule tips control mitotic spindle size

**DOI:** 10.1101/2024.02.01.578443

**Authors:** Lee-Ya Chu, Daniel Stedman, Julian Gannon, Susan Cox, Georgii Pobegalov, Maxim Molodtsov

**Affiliations:** The Francis Crick Institute, London, NW1 1AT, United Kingdom; King’s College London, London, WC2R 2LS, UK; Department of Physics and Astronomy, University College London, London, WC1E 6BT, United Kingdom

## Abstract

Mitotic spindle is a complex bipolar cellular structure that ensures chromosomes segregation between dividing cells. Correct spindle size is required for the accurate segregation and successful passing of genomes to the newly formed cells. The spindle size is believed to be controlled by mechanical forces generated by molecular motors and non-motor proteins acting in the spindle microtubule overlaps. However, how forces generated by individual proteins enable bipolar spindle organization is not well understood. Here, we developed tools to measure contributions of individual molecules to this force balance. We show that microtubule tip-trackers act synergetically at microtubule tips with minus-end directed motors to produce a system that can generate both pushing and pulling forces. We show that this system harnesses forces generated by growing tips of spindle microtubules and provides unique contribution to the force balance distinct from other force generators because it acts at microtubule tips rather than in microtubule overlaps. We show that this system alone can establish stable bipolar organization in vitro and in mitotic spindles in human cells. Our results pave the way for understanding how mechanical forces in spindles can be fine-tuned to control the fidelity of chromosome segregation.

## Introduction

During cell division, microtubules form bipolar spindles that rearrange and segregate mitotic chromosomes. Assembly of the mitotic spindle is driven by mechanical forces acting between the microtubules originating from opposite spindle poles, which determine the final size and position of the spindle. Errors in the spindle architecture lead to chromosome instability and aneuploidy, and targeting the force balance in spindles may preferentially target cancer cells while sparing normal human tissue ^1–3^. However, to identify therapeutically relevant strategies, it is important to understand how mechanical forces generated by various molecules integrate to control the spindle architecture.

Molecular motors and non-motor proteins work together to establish the size and position of the mitotic spindle ^4–6^. A key determinant of the spindle size is the balance of the pushing and pulling mechanical forces generated by the plus-end and minus-end directed motors that act between the spindle poles in antiparallel microtubule overlaps. Several motors are implicated in this balance, a combination, which is thought to ensure robustness and accuracy of the process ^7–9^. However, which forces individual molecules generate in bipolar microtubule arrays is largely unknown, and, therefore, exact roles and contributions of force generating molecules to the force balance in mitotic spindle are poorly understood.

HSET is a kinesin-14 family minus-end directed motor that supports pole focusing in mitotic spindles ^10,11^. In vitro, HSET generates pulling forces that slide antiparallel microtubule bundles in the direction opposite to the plus-end directed motors ^12,13^, which is partially mediated by the HSET tail domains that interacts with microtubules diffusively ^14,15^. This interaction also allows HSET to slide antiparallel microtubules with speeds that depend on both motor density and the size of the microtubule overlap ^16^. Activity of HSET can generate asters and microtubule networks ^17^ and it is known to affect the spindle size ^18^. However, HSET does not directly counteract plus-end motor activity that could result in efficient force balance leading to bipolar configuration ^14,19^. To the contrary, reduction of HSET causes shorter, not longer, spindles as would be expected from the contribution of a minus-end directed motor on the force balance ^8,20–22^. Consistent with this, in yeast, kinesin-14 motor analogue is required for the spindle elongation, which is also dependent on the microtubule plus-tip tracking analogue of EB ^23^, but the underlying mechanism is not understood.

Microtubule tip-tracking EB proteins autonomously recognize growing microtubule tips and act as both regulators of microtubule dynamics as well as hubs that recruit large number of other regulatory proteins to growing microtubule tips ^24^. Individual EB proteins track microtubule tips by rapid binding and unbinding to and from them. The turnover time of an individual EB dimer at the microtubule tip is approximately ∼ 1 s, while its affinity to the rest of microtubule GDP lattice is ∼10 times smaller. This ensures that collectively multiple EBs stay at the growing microtubule tips ^25–27^. EB depletion in mitosis is similar in phenotype to HSET, it causes shorter spindles ^28,29^ and may lead to misplaced and disorganized spindles as well as chromosome alignment defects resulting in chromosome segregation errors ^30,31^.

One of the most abundant elements in the mitotic spindle are the microtubules themselves. Changes in microtubule dynamics affect the mitotic spindle length ^32,33^, and even mild changes can be associated with mispositioned and disorganized spindles, chromosomal instability and aneuploidy ^2,3^. This suggests that microtubule dynamics affects the force balance in spindles, but the mechanisms underlying this dependence remain elusive.

Growing microtubule tips are also force generators. Tips of growing microtubules generate pushing forces up to 5 pN ^34^, and these forces are implicated in variety of cellular processes including rearrangement of microtubule networks ^35–39^ as well as stabilization and centring of the mitotic spindle ^40–42^. However, how forces generated by the interpolar microtubules are implicated in the assembly of spindles is unknown.

In most spindle assembly models aiming to explain the size and position of the spindle, action of individual motors is generally assumed to be independent of one another. Thus, plus-end directed, and minus-end directed motors are thought to antagonize each other in the overall balance of forces. However, recent studies show that motor density and multivalent interactions between them can influence individual motor behaviour. Both plus-end and minus-end directed motors can change their speeds or reverse directionality altogether depending on their interactions and surroundings ^16,36,43,44^.

Multivalent interactions play even bigger roles at the microtubule tips where many EB-dimers form interactions with multiple partners ^45,46^. Tails of EB dimers form interactions with the cargo binding domains of dimeric HSET. Such tail-to-tail EB-HSET arrangement exposes both molecules’ active microtubule binding domains, which enable the ensemble activity that its individual components don’t have. EB/HSET complex can reverse the minus-end directed movement of HSET to plus-end directed and guide tips of growing microtubules along the pre-existing ones facilitating organization of microtubules into parallel bundles ^36^.

Thus, multivalent interactions between molecular motors, microtubules and tip tracking proteins enhance their capabilities to rearrange microtubule networks, but also suggest that contributions of individual molecules in ensembles need to be understood within the context of their surrounding and the other molecules that they operate with. Interaction between EB and HSET must affect the force balance in bipolar organization of microtubules, but how this contributes to the maintaining size and position of the spindle is unknown. New approaches are needed to measure mechanical contributions of individual motors, non-motor proteins and microtubules in their ensembles to understand the necessary and sufficient mechanical mechanisms that allow robust and accurate spindle assembly and positioning.

Here, we developed an *in vitro* approach to study how individual components of the microtubule spindle contribute to the force balance in bipolar microtubule arrays. We discovered that forces generated by microtubules that act via tip-tracker EB and a single molecular motor HSET provide unique contribution to the force balance that is distinct from all other motors and non-motor proteins and allows for the formation of the bipolar state. We dissected the mechanism behind generation of the pushing forces by antiparallel microtubules and used computer simulations to rationalize how this system enables pole separation and achieves stable bipolar structures. Furthermore, we confirmed that contribution of forces generated by microtubule tips via EB/HSET system is essential for the correct assembly and proper size of the mitotic spindle in live human cells. Taken together, we established new framework for dissecting mechanics of the mitotic spindle assembly and identified a new class of microtubule dynamics-based force generators that provides significant contribution to the force balance in mitotic spindle.

## Results

### A single minus-end directed motor and dynamic microtubules generate stable bipolar organization

To dissect the mechanical forces that facilitate organization of the bipolar microtubule networks we established an “artificial spindle” assay based on the double bead optical trapping experiment in a multi-channel microfluidic flow cell (Figure 1A). In this assay, we attached Digoxigenin-labelled GMPCPP-stabilized microtubule seeds to two 2 μm-sized Anti-Digoxigenin coated beads, which were held in two independent optical traps. We then changed the buffer by moving the beads into a different microfluidic compartment containing 1 mM GTP, 18 μM tubulin and 10 nM CAMSAP3 to initiate the microtubule growth exclusively from the microtubule seeds. Addition of CAMSAP3 blocked the microtubule growth at the minus ends and ensured that all microtubule dynamics occurred only at their plus ends. Thus, all microtubules grew from the beads with their plus ends out mimicking the aster organization at the microtubule organizing centers.

**Figure 1.**
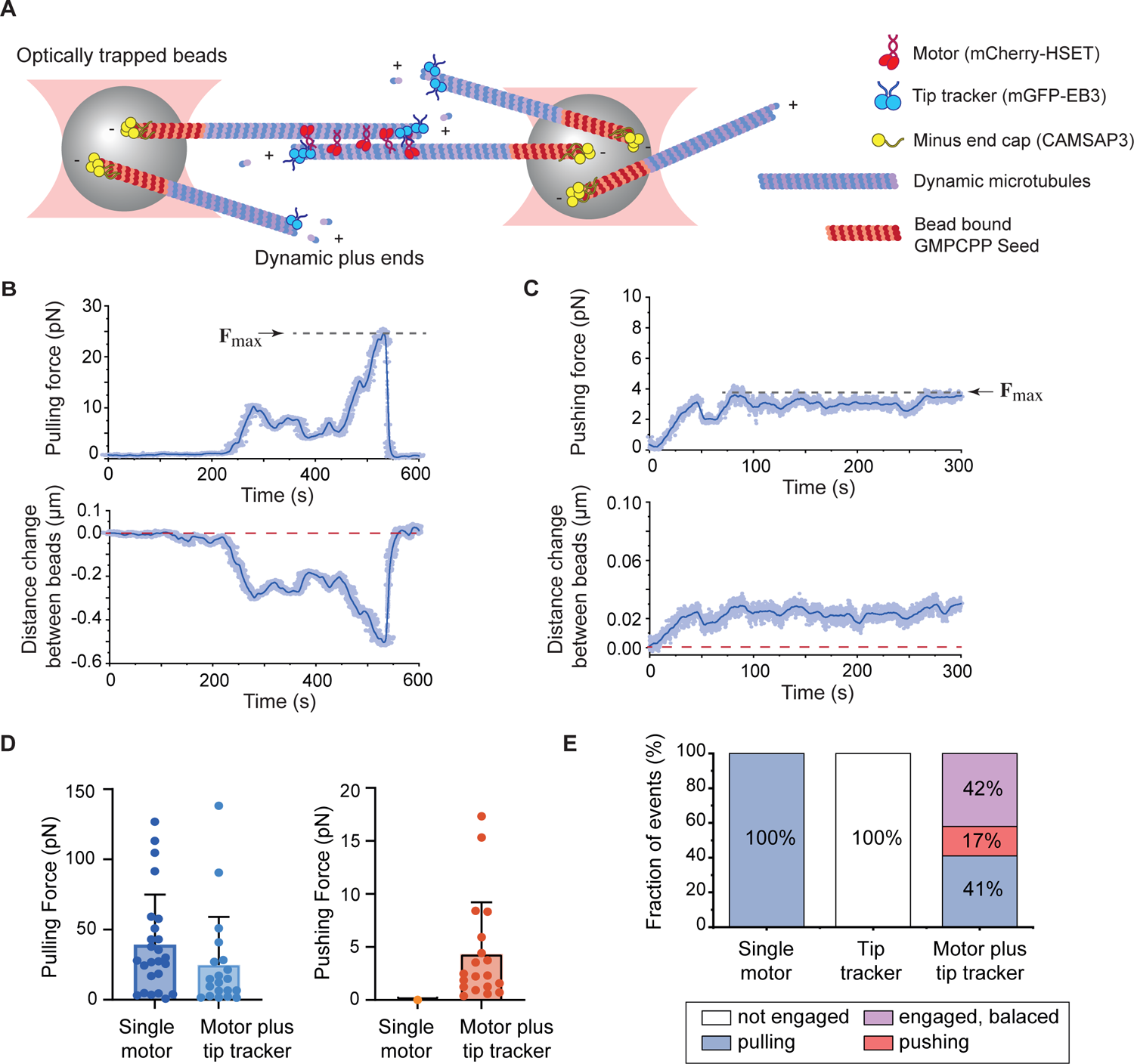
Microtubule dynamics, single molecular motor and a tip tracker stabilize bipolar organization of two microtubule asters. **A**, Schematics of the double optical trap “artificial spindle” assay. Multiple GMPCPP microtubule seeds tethered to plastic beads held in two optical traps and capped at their minus ends mimic two microtubule organization centres. The assay measures the force acting in the antiparallel microtubule overlaps. **B,** Example recording shows pulling force (top) and the corresponding decrease in distance between the beads (bottom) in the presence of HSET only. **C,** Example recording shows pushing force (top) and the increase in the distance between beads (bottom) in the presence of EB and HSET. Maximum force inferred from traces is shown in B and C. **D,** Histograms of maximum pulling and pushing forces extracted from individual traces. **E,** Quantification of the fraction of time that the system spends pushing, pulling, being balanced (with at least two antiparallel microtubules from opposite poles engaged) or disengaged (when microtubules from opposite poles do not interact) depending on the experimental conditions.

Next, we supplemented the buffer with minus-end directed motor HSET, which is known to crosslink and slide antiparallel microtubules.^12,13^ We performed our experiments with low density of microtubule seeds to minimize the number of microtubule intersections (Video S1, see Methods). At 10 nM HSET microtubules growing from the opposite beads started engaging effectively and formed antiparallel bundles. As soon as microtubules from the opposite asters engaged, forces acting on the beads began increasing, pulling them closer to one another (Figure 1B). The maximum pulling force averaged across multiple traces in this configuration was ∼ 40 pN, and in some traces exceeded 100 pN before an abrupt drop, which was either due to one of the engaged microtubules breaking or undergoing a catastrophe. The drop was typically followed by another force increase due to engagement between a different pair of other microtubules, which resulted in a saw-tooth like force pattern (see more examples in Figure S1). Thus, consistent with previous reports, we observed that HSET promoted formation of antiparallel microtubule overlaps and generated force that led to antiparallel microtubule pulling.

To complete the system, we added microtubule tip tracking protein EB3, which is known to bind HSET in a way that makes microtubule binding domains on both EB3 and HSET available for engaging microtubules.^47,48^ We verified this interaction and confirmed that it allowed HSET to transport EB away from microtubule tips on individual microtubules (Figure S2A/Video S2). Unexpectedly, however, when both minus-end directed motor and tip tracker were present in our assay, we observed not only pulling, but also pushing forces as was indicated by the increase in the distance between the beads (Figure 1C,D and Figure S1). The switching between pulling and pushing in the presence of EB3 appeared to be stochastic. To quantify this, we measured the relative amount of time the two beads experienced during pulling, pushing or while being in relative balance across all recorded traces (n = 84 traces, total recording time ∼ 14 hours). Our data showed that while microtubule engagement in the presence of HSET alone always resulted in generation of pulling forces, in the presence of both HSET and EB engagement resulted in pulling ∼ 40% of the time and pushing ∼17% of the time (Figure 1E). We also observed that in another ∼ 40% of the time microtubules growing from opposite beads were engaged in the presence of EB and HSET, but no overall force was detected indicating that pushing and pulling forces were balanced and the system was in relatively stable bipolar state (Figure 1E, Figure S1B). Finally, we verified that with EBs alone, microtubules did not engage, and we could not detect any force between microtubule asters (Figure S1C). Thus, generation of pushing forces in antiparallel microtubule networks depended on the combination of the minus-end directed motor and a tip tracker. This was an unexpected finding given that microtubule end binding proteins are non-motors, but nonetheless strongly biased the force balance in the two asters of dynamic microtubules. Even more intriguingly, our data suggested that a simple system consisting of a single molecular motor and dynamic microtubules with tip trackers could produce a stable bipolar state.

### Two growing antiparallel microtubules generate 5 pN pushing force

The maximum pushing forces that we measured in our optical trapping assay rarely exceeded 5 pN (Figure 1D). Since most of the load on the beads is likely carried out by a single pair of the antiparallel microtubules, we reasoned that this likely corresponds to the maximum force the pair can generate. However, the exact number of engaged microtubules was poorly controlled in the optical trapping assay. Thus, to verify the force generated in the EB/HSET system in a single pair of two antiparallel microtubules we employed a total internal reflection fluorescence (TIRF) microscopy assay. In this assay, we attached Digoxigenin-labelled GMPCPP stabilized microtubule seeds to the surface of Anti-Digoxigenin coated and passivated coverslip. We then added tubulin and GTP to initiate the growth on microtubule from the surface immobilized seeds as well as 10 nM HSET, and 100 nM EB. Out of all microtubules that grew from GMPCPP seeds, we selected those where two microtubule plus-ends ran into each other in approximately antiparallel configuration (Figure 2A, n = 25 total engagement events selected). After the encounter, the tips engaged in the presence of HSET, and continued to grow along one another. As expected EB formed visible comets on the tips of growing microtubules and density of HSET in the microtubule overlap was higher than on single microtubules ^16^ (Figure S2B). Notably, however, when both EB and HSET were present, microtubules that continued to grow into antiparallel overlaps started to buckle (Figure 2A, Video S3). Buckling indicates presence of pushing force acting against GMPCPP seeds as microtubules continue to elongate. Thus, similar to the optical trapping assay, buckling also indicated that EB/HSET system can use microtubule polymerization to generate pushing forces in antiparallel overlaps.

**Figure 2.**
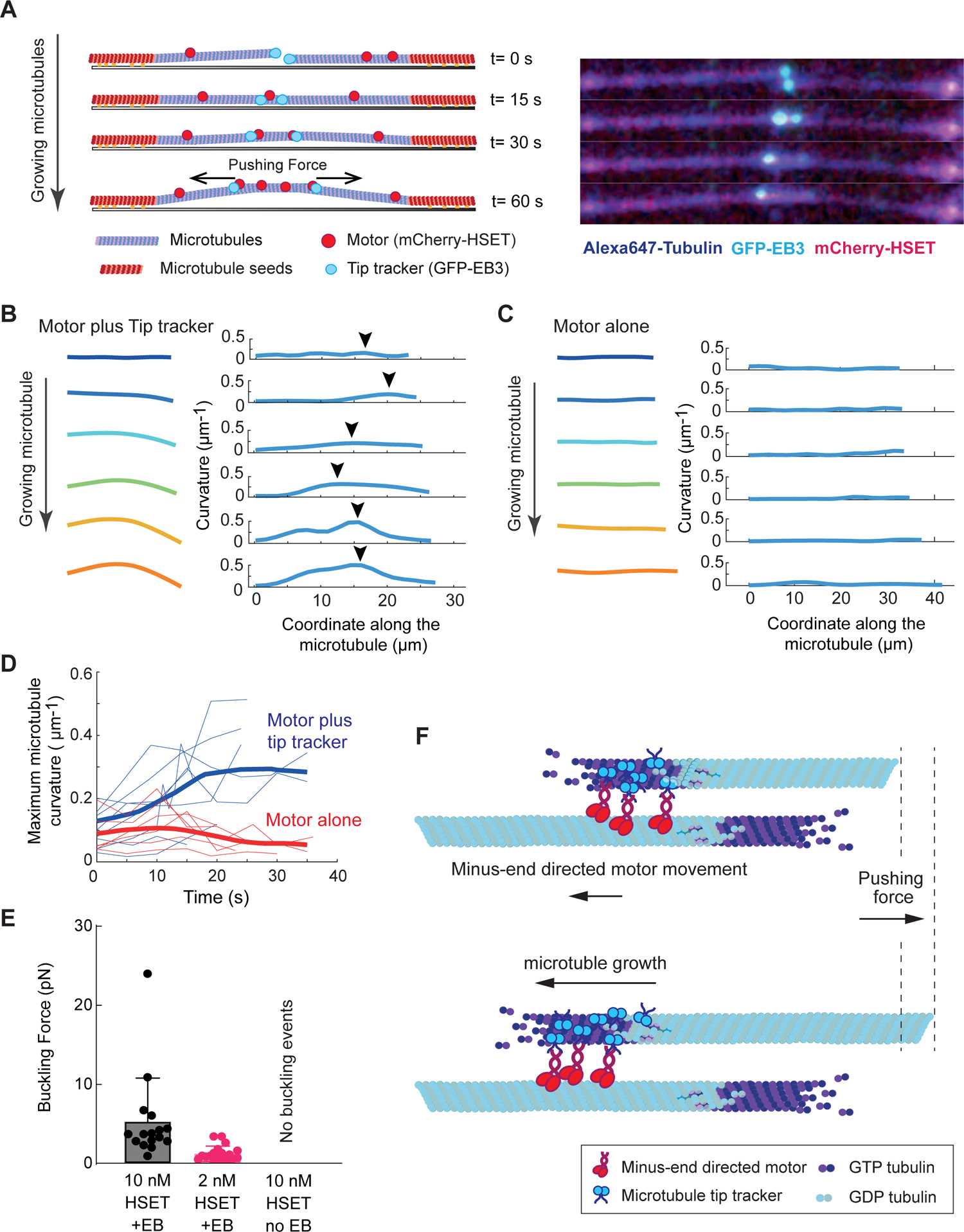
Two antiparallel growing microtubules generate pushing force. **A**, Schematics and montage images of an example buckling event between two antiparallel microtubules in the presence of EB and HSET. Alexa647 microtubules shown in blue. mCherry-labelled HSET and mGFP-labelled EB3 are red and cyan correspondingly (See Figure S2B). **B,** Shapes of a microtubule undergoing buckling in the presence of 100 nM EB3 and 10 nM HSET as it grows in antiparallel overlap inferred from images. Only one microtubule from a pair is shown. On the right - curvatures of the microtubule at the corresponding time points along the microtubule length. Arrowhead points to the maximum curvature. **C,** Same as in B, but no EB present (HSET only). **D,** Maximum microtubule curvatures determined from B and C plotted as function of time passed since the first antiparallel engagement event. Thin curves are individual examples and thick curves are averages. Blue corresponds to 100 nM EB3 and 10 nM HSET, red - 10 nM HSET. **E,** Buckling force for 100 nM EB3 plus 10 nM HSET, 100 nM EB3 plus 2 nM HSET and 10 nM HSET without EB3. **F,** Schematics of the model proposing how coupling between minus-end directed motor and a growing microtubule tip can harness pushing force generated by the microtubule growth. For simplicity protein action is shown at one microtubule tip only.

We also noticed that microtubules became visibly bent as they were allowed to grow and push. To quantify the bending process, we traced the shapes of microtubules and calculated the curvature of the microtubule along its length (Figure 2B). We then plotted the maximum curvature of microtubules as a function of time after it first engages in an antiparallel overlap and continues to grow (Figure 2B-D). This analysis confirmed that in the presence of EB and HSET antiparallel microtubule encounters are followed by microtubule buckling as indicated by the increasing maximum curvature, thus demonstrating that microtubules exert pushing force. Consistent with our optical trapping results, we did not observe detectable buckling in the absence of EBs (Figure 2C).

Growing microtubules start to buckle only when the pushing force that they generate overcomes a certain threshold, and its value can be used to infer the amount of “pushing” force that they generate.^34,49^ This “buckling force” depends on the mechanical properties of the microtubule, and can be calculated based on the microtubule length and its persistence length (see Methods). To infer pushing forces from the experiments, we collected all examples that showed buckling, and calculated the force using microtubule length extracted from their shapes. As presence of EBs was required to see the buckling, we kept constant concentration of EBs and tested how concentration of HSET affected the buckling. At 2 nM HSET concentration, we only observed buckling of long microtubules, which when converted into buckling force corresponded to forces of approximately 1pN. At 10 nM HSET buckling events became more abundant and multiple shorter microtubules buckled, indicating generation of higher pushing forces (Figure 2E). The distribution of the pushing forces was highly non-normal (p-value 4 · 10^−13^, one-sample Kolmogorov-Smirnov test) and decreased abruptly at forces above 5 pN, suggesting that this is approximately the maximum force that the system can develop. Thus, these measurements were consistent with the forces measured in the optical trapping assay and reinforced our conclusion that EB/HSET system can generate up to ∼5 pN pushing force in a single pair of antiparallel microtubules.

### Ensembles of EB dimers connected by flexible linkers move processively with microtubule tips

Next, we sought to understand how tip tracking proteins contribute to the generation of the pushing forces in antiparallel microtubule bundles. Since individual EBs bind microtubule tips only transiently, we hypothesized that the mechanism likely depends on the combined action of multiple EBs at the microtubule tip that are connected to HSET. In dividing cells, the rate of the microtubule growth is larger than the speed of HSET movement ^29,50^, which was also true in our *in vitro* experiments (Figure S2C-E). We reasoned, that if multiple EBs can bind and unbind from the microtubule tip while staying connected to the HSET moving on the antiparallel microtubule, they could possibly couple the growth of the microtubule tip to the minus-end movement of HSET along the antiparallel microtubule (Figure 2F). Since microtubule grows faster than motor proteins move, the coupling between the two would result in the overall increase of the antiparallel bundle length, which would translate into the generation of the pushing force.

The main assumption in this hypothesis is that multiple EB molecules connected to the antiparallel microtubule via HSETs can harness several piconewtons of force from the growing microtubule tip, as indicated by our double-bead optical trap and TIRF assays. In other words, we suspected that several EB dimers connected to an appropriate common mechanical scaffold could form an effective force coupling device. To test this directly, we generated ensembles containing 1, 2, 3 and 4 full length EB3 dimers covalently connected via SNAP-tag to an artificial scaffold that we made from DNA (Figure 3A). To ensure that EBs were always connected to the scaffold as dimers, we expressed recombinant EB3 heterodimers that were further stabilized by the addition of a leucine zipper and in which only one subunit had a SNAP tag that we used to attach to the DNA (Figure S3A). We then incubated SNAP-tagged EB3 heterodimers with DNA scaffolds labelled with a single Cy3 fluorophore that had 1,2,3 or 4 SNAP-ligand cites, which resulted in scaffolds with defined number of EB3 dimers (Figure S3B). We introduced these scaffolds into the flow cell with microtubules dynamically growing from the surface immobilized GMPCPP seeds and tested their ability to be driven by the tip of the growing microtubule and resist mechanical forces.

**Figure 3.**
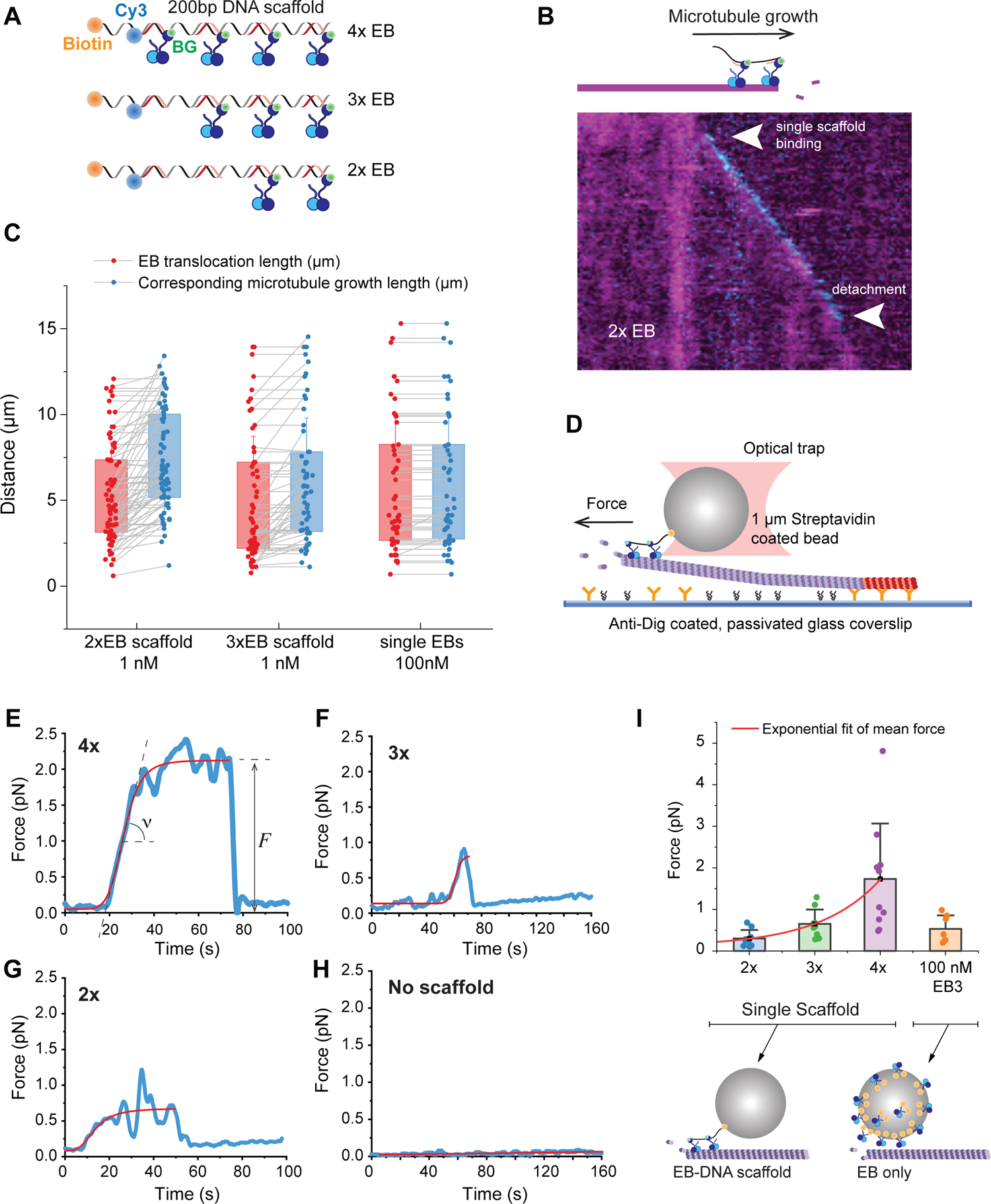
EB dimers connected by DNA scaffold move processively with microtubule tips. **A**, Schematics of EB3 ensembles on a 200 bp DNA scaffold. The DNA scaffold is based on the ∼ 200 ssDNA to which four 25 bp primers are annealed at specific locations. The primers contain SNAP-ligand for the attachment to EB3-LZ-SNAP/EB3-LZ heterodimers. The first primer is labelled with a single Cy3 fluorophore and the 200 bp DNA backbone is optionally labelled with biotin. **B,** Kymograph of a single DNA scaffold with two EB3 attached heterodimers (2x) tracking the tip of a growing microtubule. White arrows point to the scaffold binding and unbinding events to and from the growing tip. **C,** EB run lengths and MT growth distances. Red points are run lengths of the 2x and 3x single EB scaffolds (red) as well as a control where multiple single EBs (100 nM EBs) were processed by the same algorithm as scaffolds (red). Blue points are corresponding growth distances of the microtubule tips until they either underwent catastrophe or went out of the TIRF field. Grey lines connect EB scaffolds with their corresponding microtubule tips. Note that red is always below the corresponding blue as EB can only track the growing tip for the fraction of the duration of its growth, **D,** Schematics of the experimental setup for measuring forces generated by EB scaffolds. **E-H,** Individual examples of traces for ensembles with different number of EB heterodimers. Traces shown are smoothed to 20Hz and fitted with logistic function (red line) to determine the force. In E extracted force and slope are shown as *F* and ν. **I,** Combined measurements of force traces plotted as a function of the number of EBs in the scaffold. Exponential fit shows how average force increases with the number of EBs. On the right, results of the experiment in which EB dimers were coupled directly to the streptavidin bead in saturating conditions (No DNA scaffold used. One 1-μm sized bead has an estimate of >170,000 EB binding sites). In **E**-**I** 2x, 3x and 4x corresponds to single scaffolds with 2, 3 and 4 EB dimers respectively.

At 1 nM concentration, single scaffolds containing just two EB3 dimers showed processive movement with microtubule tips driven by the microtubule growth, which was illustrated by typical binding, movement and unbinding events (Figure 3B). Quantification of the fluorescent signal from the scaffold confirmed that it corresponded to a single Cy3 fluorophore and therefore a single scaffold moving with the microtubule tip (Figure S3C-F). At least two EB dimers were required for the processive movement with the growing tip. We could not detect any binding of scaffolds containing just one EB dimer or no EBs at all to the growing microtubule tips in this assay.

The processivity of the scaffolds containing just two EB3 dimers, was surprisingly high and single scaffolds could track microtubule tips for up to several microns (Figure 3C). The run length that we observed was largely limited by the microtubule dynamics and the extent to which microtubules could grow in our assay. When we increased the number of dimers from two to three, the run length did not substantially increase because it remained limited by the microtubules (Figure 3C). Note that in all cases run lengths remained below the extent of the microtubule elongation because once a microtubule starts growing it takes time for a scaffold to bind to the microtubule tip given its very low concentration in solution.

For comparison, we performed control experiments, in which we replaced 1 nM scaffolds with 100 nM GFP-labelled EB homodimers. This high concentration was required to reliably detect EB signal from EB dimers on the tips of growing microtubules because mechanism by which individual non-processive EB dimers “track” microtubule tips is different from the mechanism used by processive scaffolds containing two or more EBs. Individual EBs unbind quickly from the microtubule tips and even quicker from the microtubule lattice.^51^ High concentration of EB dimers is required to achieve high on rates needed to see the enrichment of EB dimers at the microtubule tip and compensate for the quick unbinding. Thus, we were able to distinguish two qualitatively different mechanisms of “tip tracking”. Individual EB dimers are not processive and turn over quickly at the microtubule tips, while scaffolds with several EBs can stay at the microtubule tips for much longer and move processively driven by the growing microtubule end. When we applied our processing pipeline to the data with single dimers, we found out that the signal from individual EBs at high concentration appeared on microtubule tips as soon as they began growing due to rapid binding and dissociation that occurs under high EBs concentrations. The signal disappeared with the disappearance of the growing microtubule tip. Together this yielded the same effective “run length” in this control experiments as the length of the microtubule growth (Figure 3C).

Taken together, these studies show that unlike individual dimers, EB ensembles serve as an efficient coupling device that harnesses growth of the microtubule tips to produce reliable and processive movement.

### EBs harness several piconewtons of force from single growing microtubule tips

Next, we asked how much force can be harnessed from the growing microtubule tip by the EB-DNA scaffold processively moving with it. To measure the force, we attached DNA scaffold via biotin on its end to a 1 μm size streptavidin coated plastic bead. We then trapped the bead in a single-beam optical trap, positioned it in front of the growing microtubule, and recorded the mechanical transients associated with the bead movement caused by its interaction with the growing microtubule tip (Figure 3D). We ensured that interaction between the bead and the microtubule tip was mediated by single scaffold using very low concentration of scaffolds such that only ∼20% of beads (n=42 out of 200) showed interaction (see Methods).

When growing microtubule tip exerted force on the bead, it increased, plateaued and then dropped back to zero after the growing tip passed the bead in the trap and the scaffold detached from the microtubule (Figure 3E-H). We fitted all signals with a logistic-like function to determine the speed of the force increase and the maximum force associated with the interaction between the scaffold on the bead and the growing microtubule tip. We counted only those signals in which the force increased approximately with the speed of the microtubule growth when the tip reached the bead. With only one EB heterodimer per scaffold, we failed to detect any signals satisfying these criteria. With two EB heterodimers per scaffold the detection of force generation became robust with 9 out of 50 traces showing the increase in force with the speed of microtubule growth and the maximum force of ∼ 1 pN (Figure 3G).

When we tested scaffolds with larger number of EB dimers, we found that the maximum force increased approximately exponentially with the number of EB dimers in a scaffold (Figure 3I). The maximum number of EB dimers per scaffold that we could test was four, but even with this relatively low amount, forces frequently reached above 2 pN (average force was 1.7 pN and standard deviation 1.3 pN), which is a significant contribution to the maximum force of 5 pN that a single microtubule tip can generate. Given the high variability of the measured forces and assuming the exponential dependence inferred from our data, we extrapolate that possibly as little as five to eight EB dimers could already be sufficient to harness pushing forces up to 5 pN from a growing microtubule tip. In a control experiment, we coated streptavidin beads directly with saturating amount of biotinylated EB3 dimers. In this case, up to a hundred of thousands of EBs are expected to be attached to the bead, but we could only detect forces below 1 pN (Figure 3I, 0.5 ± 0.3 pN mean ± standard deviation), which were consistent with previous measurements.^38^ Thus, connection by a flexible scaffold enhances ability of EBs to harness mechanical force from the growing microtubule tips.

### Microtubule/EB/HSET system generates dynamic bipolar microtubule organization

To gain further insight into the force generating properties of EB/HSET system and understand how it could facilitate stabilization of the bipolar configuration, we used the open source software package Cytosim.^52^ We simulated the interaction between two 2D asters of microtubules in the presence of HSET motors and EB. Microtubules were modelled as elastic rods with their minus fixed at one of the two microtubule nucleation centers, and dynamic plus ends. The HSET and EB/HSET were both modelled as motors consisting of two units connected by an elastic linker (Figure 4A,B). In the case of HSET, both units stochastically bound the microtubule lattice and moved towards the minus ends of the microtubules. For the EB/HSET motors, the unit representing HSET bound the microtubule lattice and moved towards the minus end. The unit representing the EB bound and tracked the microtubule tip and could resist external forces up to 5 pN as suggested by our experiments. Thus, in this simplified model, the EB domain of the EB/HSET system represented the collective action of multiple EBs that remained bound at the microtubule tip as a scaffold stabilized by their interaction with the other microtubule via HSET (Figure 4C).

**Figure 4.**
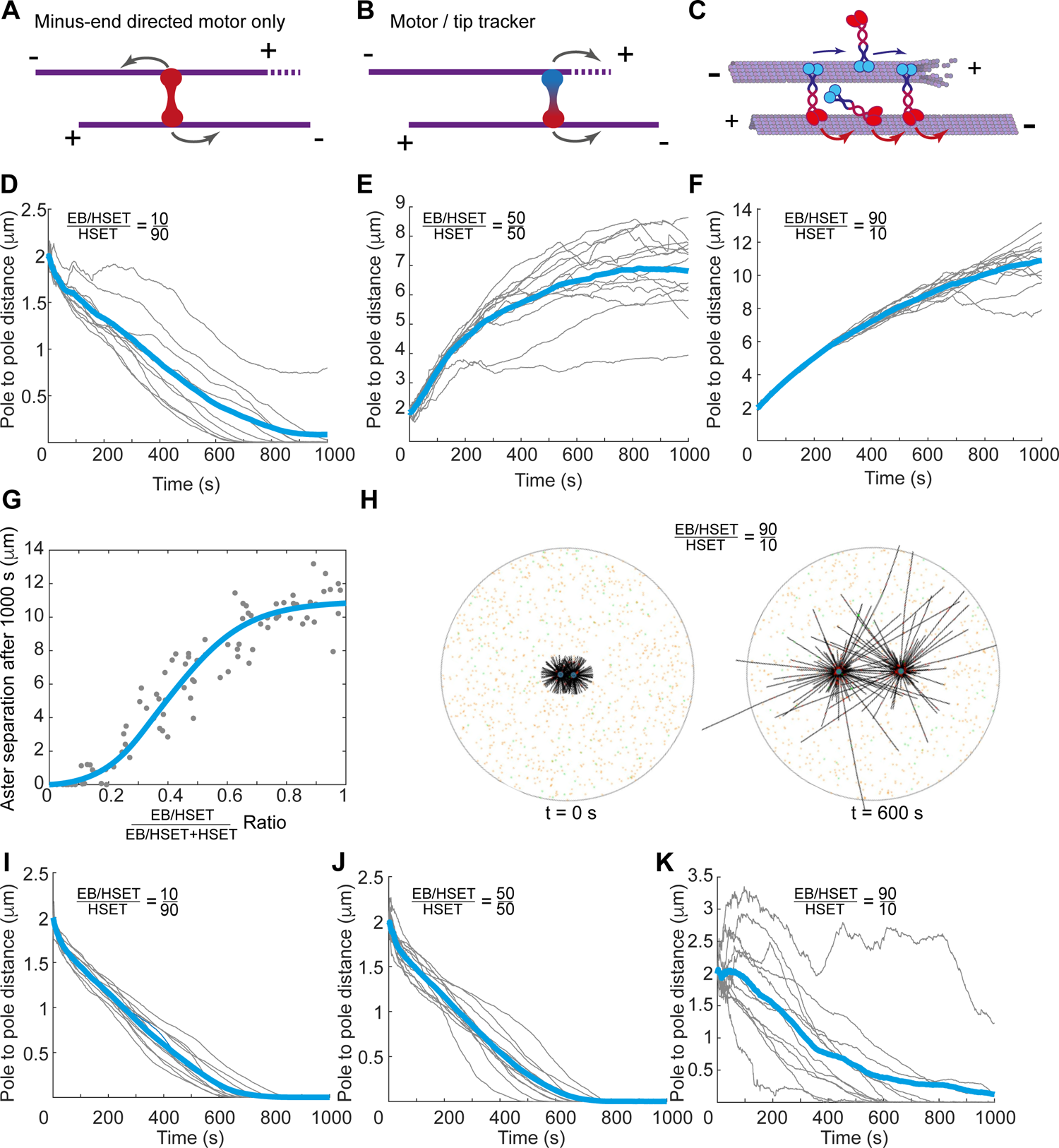
Organization of the two microtubule asters by the microtubule/EB/HSET system. **A**, Schematics of the activity of minus-end directed HSET motors in an antiparallel microtubule overlap. **B,** Schematics of the activity of the tip tracker/motor (EB/HSET) system in an antiparallel microtubule overlap. **C,** Multiple EBs connected to HSET on antiparallel microtubule represent one EB/HSET simulation unit. **D-F,** Pole-to-pole distances as a function of time in the simulations for different ratios between HSET and EB/HSET (shown on top). Grey lines are individual simulations, thick blue lines – average between them. Microtubule growth was assumed 2x faster than HSET movement (as in Fig. S2B). **G,** Pole-to-pole separation distance after 1000 s of simulation as a function of the relative fraction of the EB/HSET in the system. Dots are individual simulations, line is a trendline only. **H,** Snapshot images of the simulations for the 90% EB/HSET system in the beginning and in the middle of the simulation (Corresponds to the Movie S6). Colours: Black – microtubules, Green – unbound HSET, Orange – unbound EB/HSET, Dark green – microtubule bound HSET, Red – microtubule bound EB/HSET. **I-K,** Pole-to-pole distances as a function of time in the simulations for different ratios between HSET and EB/HSET in which EB/HSET was assumed to detach at forces exceeding 0.1 pN. Notations follow **D-F**.

We assumed the fixed amount of HSET and EBs in the simulated volume with two microtubule asters, and we varied the ratio between HSET and EB/HSET complexes. As expected, when the interaction between microtubules was dominated by HSET alone, antiparallel microtubules were pulled together, and the two asters rapidly fused (Figure 4D, Video S4). However, when the fraction of HSET/EB was increased, the pushing force prevailed at the initial stages of the aster separation and then plateaued leading to stable bipolar configuration (Figure 4E,F, Video S5). The steady-state distance between the asters did not depend on the absolute number of the molecules, and decreasing the absolute amount of both HSET and EB/HSET 20-fold resulted in a similar steady state aster-to-aster distance albeit with higher fluctuations due to smaller overall number of motors (Figure S4A). The distance did depend on the microtubule dynamic instability parameters and increasing the microtubule growth rate resulted in the increase of the average distance asters (Figure S4B).

These simulations confirmed that when EBs at the tip of the microtubules provide connections to HSET motors that can resist ∼ 5 pN forces in antiparallel overlaps, the EB/HSET system can stabilize bipolar configuration of the two microtubule asters. To further bolster this conclusion, we compared these simulations to the simulations in which we assumed that EBs cannot transmit force from microtubule tips and detach at forces exceeding 0.1 pN as would be expected for individual unconnected EB dimers. Indeed, although these EBs can rebind to microtubule tips in the absence of load, our simulations revealed that in this case asters fused and could not maintain bipolar configuration at any of the EB/HSET to HSET ratios (Figure 4I-K). Thus, ability of EB to harness forces generated by microtubule growth enables EB/HSET system generate pushing forces between the microtubule asters and may facilitate establishment of the bipolar configuration.

### Pushing forces generated by antiparallel microtubules are required for correct spindle size in human cells

To determine the contribution of the pushing forces generated by the microtubule/EB/HSET system to the force balance in the metaphase mitotic cells, we chose H1299 non-small-cell lung cancer cells with π-EB1 system.^53^ In these cells, endogenous EB proteins are removed and replaced with light-controllable π-EB1. In the absence of blue light these cells have fully functioning EB. However, upon blue-light activation, the plus-end tip (+TIP) binding domain of the π-EB1 dissociates from the microtubule, which abolishes the interactions between π-EB1 and all its +TIP binding partners including presumably HSET (Video S6). Thus, all endogenous EBs in these cells are rapidly inactivated by blue light.

To dissect the specific contribution of the EB/HSET interaction to the force balance in mitotic spindles, in addition to π-EB1 system, we introduced another pair of EB and HSET molecules, whose interaction we could control independently (Figure 5A). To develop this system, we generated EB1 in which C-terminal +TIP binding domain was replaced with FRB (EB1ΔC-mApple-FRB), and an HSET in which the tail region containing cargo and EB binding domain was truncated and replaced with an FKBP (FKBP-mGFP-HSETΔN). Thus, we could control the interaction between the EB1ΔC-mApple-FRB and FKBP-mGFP-HSETΔN by adding rapamycin. Without rapamycin these molecules did not contribute to the spindle size and organization as both lacked their protein interaction domains and therefore did not interact with each other or other partners. Rapamycin addition led to the formation of the EB/HSET complex and allowed us to determine the effect of the presence of this complex on the dynamics of the spindle in the presence or absence of other EB interactions controlled by blue light.

**Figure 5.**
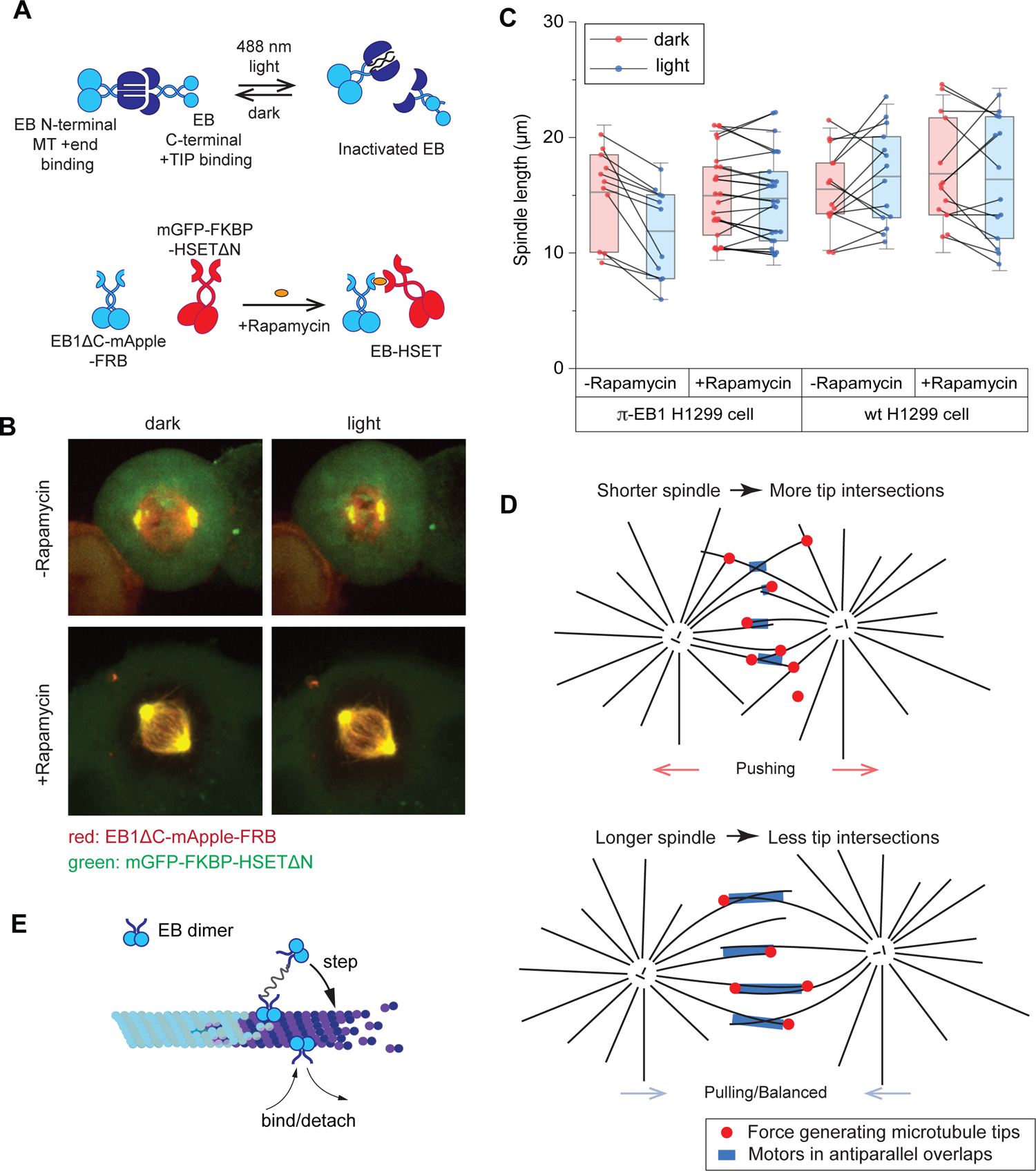
Metaphase spindle length in H1299 cells depends on the pushing force generated by EB/HSET. **A**, Schematics of the two systems that allow endogenous EB light inactivation and rapamycin induced EB – HSET interaction. **B,** Fluorescent images of π-EB1 H1299 cells with or without rapamycin treatment, light and dark blue-light activation. EB light inactivation induces spindle shortening, but only in cells lacking additional rapamycin activated EB/HSET system. **C,** Spindle length measured in H1299 and π-EB1 H1299 cells before (red points) and after (blue points) EB light deactivation with and without rapamycin. Grey lines connect single cell spindle length measurements before and after light activation. The mean spindle length for π-EB1 H1299 cells are - Rapamycin, dark: 15 ± 4 µm; - Rapamycin, light: 12 ± 4; +Rapamycin, dark: 15.0 ± 3.7 µm; +Rapamycin, light: 14.7 ± 3.9 µm. The mean spindle length for wild-type H1299 cells are - Rapamycin, dark: 15.5 ± 3.5 µm; - Rapamycin, light: 16.6 ± 4.2; +Rapamycin, dark: 17 ± 4.6 µm; +Rapamycin, light: 16.4 ± 5.3 µm (all numbers are mean ± SD). **D,** Proposed model how a single minus-end directed motor and pushing microtubule tips separate microtubule asters and stabilize bipolar organization. See text for details. **E,** Schematics of how flexible connection between two microtubule tip-tracking dimers enables their processive movement with the growing microtubule tip.

Without rapamycin treatment, mitotic spindles shortened by ∼23% in π-EB1 H1299 cells expressing FRB-mApple-EB1C and FKBP-mGFP-HSETΔN after exposure to blue light (Figure 5B,C, Video S7). This was consistent with what has been measured for these cells in the absence of the additional FRB-mApple-EB1C and FKBP-mGFP-HSETΔN ^29^.

This also indicated that EB/HSET interaction may have provided pushing forces in the spindle, and when these forces were removed, the new balance of forces resulted in a shorter spindle length. To test this further, we treated cells with rapamycin before exposing them to the blue light. In this experiment, we observed that the spindle length increased ∼20% comparing to non-rapamycin treated cells under blue-light stimulation (Figure 5B,C, Video S8). Interestingly, the spindle length in this case was indistinguishable from the normal spindle length in cells without blue illuminations and a wild-type EB/HSET system (Figure 5C). Moreover, rapamycin treatment made the spindle size insensitive to the blue light exposure and consistent with the spindle length in wild type cells.

Thus EB/HSET interaction changes the force balance in spindles leading to the increase of the spindle length. This is consistent with our *in vitro* data showing that microtubule/EB/HSET system generates pushing forces in the antiparallel microtubule overlaps and demonstrates that this system facilitates establishment of the spindle size in metaphase cells.

## Discussion

In this work, we developed approaches to directly measure mechanical contributions of individual molecules to the force balance in bipolar microtubule arrays. We showed that a simple system consisting of a single minus-end directed motor that acts synergetically with growing dynamic microtubules via the action of tip tracking EB proteins can perform both, separation of the two microtubule asters and stabilization of their bipolar organization. HSET is known to affect the aster formation, the spindle size and to interact with EB proteins ^15,17,18,48^, and EBs are known to make multivalent condensates and facilitate transport ^38,39,45,46^. Here, we reveal how the combined EB/HSET/microtubule system allows harnessing microtubule forces that stabilize the bipolar organization of two asters, which is not achievable by mechanisms used by these molecules alone. This represents a new, microtubule tips-driven force generation mechanism, which plays critical role in regulating spindle size in metaphase cells.

Compared to the force generated by motor and non-motor proteins, the microtubule tip generated force in the mitotic spindle exhibits two major differences making this force generation mechanism distinct and unique. First, for motors and crosslinkers the total force that they exert in a microtubule overlap depends on the overall size of the overlap, with larger overlaps generating more force. However, forces generated by the microtubule/EB/HSET system do not depend on the size of the overlap but depend on the density of the microtubule tips and the probability of the intersection between the tip and the lattice of the opposite pole microtubule. This explains how this system alone can build and stabilize bipolar spindle.

When two asters are in proximity and the size of the spindle is small, the density of the microtubule tips is high and pushing forces that they generated dominate over forces generated in the microtubule overlaps (Figure 5D). As poles separate, density of the intersection between microtubule tips and lattices drops, which decreases the pushing force and leads to a balanced bipolar state.

The second important difference is that unlike all other motors, multiple EB/HSET molecules form a complex at microtubule tips that effectively combines the minus-end and plus-end directed activities in one system. Such hypothetical motors were theoretically predicted to be able to generate stable bipolar configurations,^54^ but experimental evidence for their existence has been lacking. Our data present first evidence that they may indeed exist and be formed by combined action of separate molecules: while HSET moves to the minus-end, EBs on the tips of the growing microtubule end move towards the plus-end direction. Thus, the minus-end directed action of HSET drives tips of microtubules towards the opposite pole, but only for growing microtubules. The rate of microtubule growth is always faster than the movement of HSET in both *in vitro* and *in vivo* resulting in the generation of dynamically maintained antiparallel overlaps. This creates conditions for the stabilization of the bipolar architecture.

Our study also revealed the mechanism that EB proteins use to harness the force generated by microtubule growth. The data showed that just two EB dimers connected by a flexible scaffold become an efficient coupling device that can be driven by the growing microtubule tip with remarkable processivity. This was unexpected because individual EB molecules turnover at microtubule tips with a lifetime of ∼ 1s,^55,56^ and EB proteins are not motors themselves. We also discovered that when connected by a flexible scaffold, force generated by EBs increases exponentially with the number of EB dimers in the scaffold when the number of dimers is small. Interpolation to the higher number of EBs showed that possibly less than ten EB dimers might be sufficient to harness 5 pN of force, which is approximately the maximum pushing force generated by EB/HSET system that we measured per single antiparallel microtubule pair.

The maximum force harnessed by EB/HSET from the growing microtubule tip is remarkably similar to the maximum force that a growing microtubule can generate by pushing against a barrier.^34^ This is surprising given the two molecular mechanisms used for the force generation are different. However, the upper limit of the force generation in both cases is set by the same thermodynamic constraint. Maximum force that a system can generate is the energy input divided by the distance over which this energy is expended. In both cases, chemical energy is provided during attachments of the new tubulins to the microtubule tip.

The distance is also determined by the microtubule growth and therefore it is also the same, suggesting that the expected maximum generated force should be equal in both mechanisms. Importantly, the upper limit of five piconewtons suggests that increasing the number of EB dimers in the scaffolds beyond eight to ten would not increase the force further. Therefore, given there can be many dozens of EB dimers at the microtubule tips during growth, we expect that only a small fraction of them would be involved in productive force generation in complexes with HSET.

The geometry and flexibility of the scaffold are most likely the key factors that enable EBs to harness close to 100% of the force generated by a microtubule tip. The flexibility presumably allows EB dimers that dissociate from the microtubule to diffuse locally, reach and bind to new positions closer to the microtubule tip, while others keep the scaffold tethered to the microtubule surface (Figure 5E). The reason that the maximum force is reached in the presence of less than ten EB dimers is likely related to the geometry of the scaffold and the structural requirement for the transition between GDP lattice and the lattice recognized by EBs. The scaffolds only move directionally in the limited area where one lattice configuration transitions into another. EBs that dissociate in this area will have preference to bind close to the tip and therefore move directionally, while EBs positioned away from this boundary would have no preferred direction and therefore would not contribute to the force generation. The size of this transition area is likely less than 150 nm,^57^ which is consistent with our estimate that increasing the size of the scaffold above ∼ 10 EB dimers will likely push them beyond this area, thus significantly decreasing the ability of the scaffold to harness microtubule generated force.

In our experiments we used DNA as an artificial scaffold to investigate the effect of mechanical coupling between EBs on their ability to harness force generated by microtubule polymerization. Although DNA scaffold is very different in nature from scaffolding made by HSET, it presumably allows for the similar mechanical flexibility of the EB dimers and enables their efficient force transduction: in the DNA scaffold, neighbouring EBs are separated by 23 bases of the single-stranded DNA and 25 bases of the double-stranded DNA, which results in the total distance of up to ∼ 23 nm.^58^ This is in good agreement with the length of the coiled neck in HSET (∼ 25 nm) and the distance between neighbour HSETs defined by the microtubule lattice (Figure S5). Therefore, both types of connection should allow for similar EB flexibility. Thus, although we could not control or determine the exact number of EBs in the experiments with HSET, we expect that they harness forces generated by microtubule polymerization using the same mechanism as when placed on artificial DNA scaffolds given the similarity of the overall geometry of the connection.

Next, we asked whether the mechanism we discovered is required for the correct assembly of the mitotic spindles in human cells. We observed that mitotic spindles shorten ∼ 20% following rapid inactivation of EBs in metaphase and this shortening is rescued by reinstating the interaction between EB and HSET. Previously, disruption of the force balance and spindle shortening after EB inactivation was attributed to the EBs being required to engage cortical pulling forces in metaphase.^29^ However, this decrease in pulling on astral microtubules can work together with the decrease in pushing on the antiparallel spindle microtubules. Our results show that EB/HSET system generates such pushing forces in metaphase spindles as its addition restores longer spindles. Interestingly, the length of the spindle is rescued completely suggesting that EBs may perform their major function in mitotic spindles in the complex with HSET as was suggested in yeast.^23^ This also shows how HSET functions to generate pushing, not pulling, forces in mitotic spindles, which explains why HSET deletion leads to shorter mitotic spindles.

The finding that spindles shorten only 20% in metaphase suggests that after EB deactivation the system finds its new equilibrium with other mechanisms that prevent further shortening. This equilibrium could result from the action of other motors and non-motor crosslinkers that stabilize antiparallel overlaps at this stage. We speculate that pushing forces generated by microtubule/EB/HSET system plays even more important role at the earlier stages of the spindle formation before metaphase when pushing forces are required for the spindle poles’ separation, and the emerging antiparallel microtubule overlaps are not yet well stabilized by other molecules. This is supported by our simulations, which show that EB/HSET system can lead to the separation of two microtubule asters and stabilization of the bipolar configuration, which in live cells is likely then stabilized further by action of additional mechanisms.

Lastly, our findings suggest a possible explanation of how relatively mild changes in the microtubule dynamics can lead to increased chromosome instability. Alterations in the spindle architecture are mediated by the change of the force balance and therefore microtubule dynamics needs to affect the force balance to exert effect on the spindle shape. We have shown that microtubule generated force directly contributes to the assembly of the mitotic spindle through a single molecular motor and a tip-tracker, and that this system alone can provide bipolar organization of microtubule asters. Thus, microtubule generated force likely contributes to the fidelity and robustness of the spindle assembly and positioning, and, by extension, changes in microtubule dynamics would have a major impact on mitotic chromosome segregation and stability.

## Methods

### Cloning, expression and purification of SNAP-LZ-EB3 heterodimer

The DNA sequence of parallel Leucine zipper pair GCN4_v4 (LZ, IASRMKQLEDKVEELLSKNYHLENEVARLKKLVGECEGL) and GCN4_v4 with SNAP tag (LZ-SNAP) were customised by GeneArt (Invitrogen) into pMA plasmids. DNA sequence of EB3 were obtained from pETMz mGFP-EB3 (gift from Thomas Surrey lab) were cloned by infusion (Takara Bio) into the GCN4_v4 plasmids using the primer (EB3_LZ_for: 5’-GTATTTTCAGGGCGCCATGGCCGTCAATGTGTACTCCACATCTG-3’, EB3_LZ_rev: 5’-CACCGCCACCGGCGCCGTACTCGTCCTGGTCTTCTTGTTG-3’). The assembled EB3-LZ constructs were then subsequently cloned into a pETDuet-1 plasmid using restriction digestion and ligation. The EB3-LZ-SNAP were inserted into the first multiple cloning site containing N-terminus His-tag using the restriction enzymes (BamHI and SalI). The EB3-LZ were inserted into the second multiple cloning site containing Strep-tag using the restriction enzymes (NcoI and XhoI). The expected molecular weight for the EB3 heterodimers is ∼93kDa (EB3-LZ-SNAP: 56.24 kDa and EB3-LZ: 37.10 kDa).

The plasmid for co-expression was transformed into BL21 (DE3) pLyS cells. Single colonies were grown overnight for 12 hours and picked from Ampicillin resistant LB plates. Bacteria cells were inoculated into 50 ml culture and later transferred to 4L culture at 37°C in 100 μg/ml Ampicillin LB media until the O.D.600 reached 0.6. The *E. coli* culture were then induced with 0.8 mM IPTG for 12 hours overnight at 18 °C. The induced culture was then harvested by centrifuging at 3500 rpm for 30 min in a Beckman Coulter Avanti J-25 ultracentrifuge using a JLA-8.1 rotor (Beckman Coulter). The pelleted cells were resuspended in 150 ml of Lysis buffer (50 mM sodium phosphate buffer, pH 7.5, 400 mM KCl, 5 mM MgCl_2_, 0.5 mM β-mercaptoethanol) supplemented with Complete EDTA free protease inhibitor (Sigma). The cells were lysed using an microfluidizer (Microfluidics M-110L) and the lysate were cleared by centrifugation at 4 °C, 45,000 rpm for 30 min, in a Beckman Coulter Optima L-100XP ultracentrifuge using a Ti45 rotor (Beckman Coulter).

The cleared lysate was loaded onto a HiTrap 1ml Chelating column (Cytiva) charged with Nickel chloride using an ÄKTA pure™ 25 (Cytiva) protein purification system. The column was washed with Nickel buffer A (50 mM sodium phosphate buffer, pH 7.5, 400 mM KCl, 5 mM MgCl_2_, 5 mM Imidazole, 0.5 mM β-mercaptoethanol) for 2 column volumes and eluted with Nickel buffer B (50 mM sodium phosphate buffer, pH 7.5, 400 mM KCl, 5 mM MgCl_2_, 500 mM Imidazole, 0.5 mM β-mercaptoethanol) for 20 column volumes. Peak fractions were collected and pooled together, then loaded onto a Strep-Trap 5 ml column (Cytiva), column was washed for 2 column volume with Strep Buffer A (50 mM sodium phosphate buffer, pH 7.5, 300 mM KCl, 5 mM MgCl_2_, 0.5 mM β-mercaptoethanol) and eluted with a gradient of Strep buffer B (50 mM sodium phosphate buffer, pH 7.5, 300 mM KCl, 5 mM MgCl_2_, 2.5 mM desthiobiotin, 0.5 mM β-mercaptoethanol) for 20 column volumes. Peak fractions were collected and pooled together, buffer exchanged into the Lysis buffer without protease inhibitor using a disposable PD-10 desalt column (Cytiva), added TEV protease and incubated for 12 hours overnight at 4 °C. After TEV protease cleavage of the His- and Strep-tag, the sample were passed through the Nickel column and Strep column again to remove the TEV protease and protein tags. The final protein samples were pooled and concentrated to 500 μl using Vivaspin® 20 Ultrafiltration Unit with a 30 kDa molecular weight cut-off. The sample was passed through size exclusion Superdex 200 column (Cytiva) in size exclusion buffer (50 mM sodium phosphate buffer, pH 7.5, 300 mM KCl, 5 mM MgCl_2_, 0.5 mM β-mercaptoethanol), and the peak fractions were collected and pooled, the final concentration of 1 mg/ml SNAP-LZ-EB3 were aliquoted and flash frozen in Liquid Nitrogen and stored in −80°C.

### Synthesis of DNA scaffold backbone for EB3 ensembles

DNA strands were purchased from Merck. DNA backbone was generated by splint ligation of the two sequences: bb_fragement_1: 5′-AAGACACCTGAGGACTGTACCTATTTTGCGGCGAGAGGGACGACAGAAGGCTAATGTGGTGC-3′, and bb_fragment_2: 5′-[Phosphate]AGCATGATACGCGCAGGGGTCAATCGAAATGAGCGGAACCGGGGAAT TGTAACTACTCCTAGACCAACGGATGCTTGTTTACGTGCCCATATCTATAAGCTGGATCACTTCAGTTCGGCCAAGAA-3′, using the splint_bb1_bb2: GTATCATGCTGCACCACATT. Splint ligation reaction was done by mixing DNA in equal molar ratios of 1 mM in a 20 μl reaction and annealed in 1✕ ligation buffer (NEB) over 4 hours using a thermocycler (bio-rad). After ligation, reactions were diluted 100 times with PCR grade water (Thermo) and 45μl of the diluted product was supplemented with 5μl of 10✕ ligation buffer and 1 μl of T4 DNA ligase and incubated at 16 °C overnight. The ligated 187 base product was separated on a 15% TBE-UREA denaturing gel and the ligated DNA band was cut out and electro-eluted from the gel piece. For the optical trapping experiments, we used modified sequence, which contained biotin: bb_fragment_2_biotin: 5′-[Phosphate]AGCATGATACGCGCAGGGGTCAATCGAAATGAGCGGAACCGGGGAAT TGTAACTACTCCTAGACCAACGGATGCTTGTTTACGTGCCCATATCTATAAGCTGGATCACTTCAGTTCGGCCAAGAA[Biotin TEG]-3′. This was used for generating DNA backbone instead of bb_fragment_2.

Amino-labelled and biotin-labelled 25 base DNA complementary strands (Cy3R1: 5’-[Cy3] CCAGCTTATAGATATGGGCACGTAA[AmC6dT]T-3’, R2: 5’-TAGTTACAATTCCCCGGTTCCGCTC [AmC6dT]T-3’, R3: 5’-TCATGCTGCACCACATTAGCCTTC[AmC6dT]T-3’, R4: 5’-TAGGTACAGTCCTCAGGTGTCTT[AmC6dT]T) were coupled with SNAP-substrate NHS ester (BG-GLA-NHS, NEB). The DNA strands at 1 mM concentration were incubated with 10 mM of BG-GLA-NHS and reactions were cleaned up using a Bio-Spin P-30 column (Bio-rad).

Depending on the required number of the EB heterodimers in the scaffold only specific strands were coupled to BG-GLA-NHS. For example, to assemble scaffolds with two EB heterodimers, R3 and R4 were coupled to BG-GLA-NHS and Cy3R1 and R2 were left unmodified.

The 187 base DNA backbone was then mixed with all four R1-R4 primers (some of which were modified to carry Snap ligand depending on the desired scaffold) at equal molar ratio and annealed in anneal buffer (10mM Hepes, pH7.5, 50 mM NaCl, 1mM EDTA) for 4 hours using a Thermocycler (Bio-rad). The annealed products were separated on an 8% Native TBE gel and the target bands were cut out and purified by electro-elution. The assembled scaffold was concentrated using ethanol precipitation to 1 μM. To couple EB heterodimers, the scaffold was incubated with 6 μM SNAP-LZ-EB3/LZ-EB3 heterodimers at 37°C for 30 min in reaction buffer (20 mM KoAC pH 5.2, 100 mM NaCl, 1 mM DTT) then quickly moved on ice. The reactions were then aliquoted and flash frozen, and stored in −80 °C.

To verify EB3 heterodimer coupling to the scaffold we did SDS polyacrylamide gel. However, we did not heat the sample in order to preserve the structure of the DNA scaffold. Presence of SDS was sufficient to break EB3 heterodimers into monomers, but it doesn’t denature dsDNA. Boiling the sample would otherwise detach heterodimers from the scaffold. Smaller weight bends are seen on the gel correspond to scaffolds with less EBs. However, the major band always corresponded to scaffolds with expected number of attached EB heterodimers. Presence of a small fraction of scaffolds with smaller number of EBs than expected did not affect interpretation of the experiments.

### Expression and purification of mGFP-EB3, mCherry-HSET, mGFP-CAMSAP3

Expression and purification of mGFP-EB3 homodimers was done using the same protocol as for SNAP-LZ-EB3 heterodimer, while omitting the Strep column purification step. Expression of mCherry-HSET was done using the Bac-to-Bac insect cell expression system according to the manufacturer’s protocol (Thermo). Cell pellets from 500 ml of cell culture expressing Strep-tagged mCherry-HSET were resuspended in 40 ml of Lysis buffer (50 mM sodium phosphate pH 7.5, 300 mM KCl, 5 mM MgCl_2_, 0.5 mM ATP, 1 mM EGTA and 5 mM β-Mercaptoethanol). Cells were homogenised by douncing 40 strokes on ice using a dounce homogenizer and the lysate were spun for 45 minutes at 50,000 rpm, 4 °C in a Beckman Coulter Optima L-100XP ultracentrifuge using a Ti70 rotor. The cleared lysate was loaded onto a StrepTrap 5 ml column (Cytiva) and washed for 2 column volumes of Lysis buffer (50 mM sodium phosphate pH 7.5, 300 mM KCl, 5 mM MgCl_2_, 0.5 mM ATP, 1 mM EGTA and 5 mM β-Mercaptoethanol), then eluted with a gradient of Elution buffer (50 mM sodium phosphate pH 7.5, 300 mM KCl, 5 mM MgCl_2_, 0.5 mM ATP, 1 mM EGTA, 2.5 mM desthiobiotin and 5 mM β-Mercaptoethanol) for 10 column volumes. Peak fractions were pooled together and incubated with TEV protease overnight at 4 °C. The sample was passed through the Strep-Trap column again and flow through was collected and concentrated down to 500 μl before the size exclusion chromatography. Sample were injected into size exclusion column Superose 6 10/300 (Cytiva) equilibrated with Size exclusion buffer (50 mM sodium phosphate, pH7.5, 300 mM KCl, 1 mM MgCl_2_, 0.1 mM ATP, 1 mM EGTA and 5 mM β-Mercaptoethanol) peak fractions were pooled and concentrated to 500 μl before flash frozen and stored in −80 °C.

Strep-tagged mGFP-CAMSAP-3 were expressed in HEK293T cell lines, mammalian expression plasmid for mGFP-CAMSAP-3 was a gift from Dr. Anna Akmanova’s lab. HEK293T cells were seeded in Dulbecco’s Modified Eagle’s Medium (Gibco) supplemented with 10% FBS (Gibco) and Pen/Strep (Gibco), at 20 % confluency in four 15 cm tissue culture dishes (Corning). Cells were transfected using lipofectamine 3000 reagent (Thermo) according to the manufacturer’s protocol. Growth media was exchanged after 24 hours and cells were checked for protein expression using a fluorescent cell imager (Bio-Rad). Cells were harvested after another 24 hours in cold PBS before purification. Purification were done by resuspending the cells in 2 ml of lysis buffer (50 mM HEPES, pH 7.4, 300 mM NaCl, 0.5% Triton X-100, 1 mM MgCl_2_, 0.1 mM ATP and 1 mM EGTA) supplemented with Complete EDTA free protease inhibitor (Sigma) and incubated on ice for 15 min. Lysate were then spun at 14,500 rpm for 20 min at 4°C. Supernatant were collected and mixed with Strep-Tactin sepharose (IBA-Lifesciences) washed with lysis buffer and eluted with elution buffer (50 mM HEPES, pH 7.4, 150 mM NaCl, 1 mM MgCl_2_, 1 mM EGTA, 0.05% Triton X-100, 2.5 mM desthiobiotin, 0.1 mM ATP, and 2.5 mM DTT). The eluted samples were aliquoted and flash frozen and stored in −80 °C.

### Generating DIG-labelled GMPCPP stabilised microtubule seeds

Digoxigenin labelled GMPCPP microtubule seeds were made by mixing 12 μM of 15% Digoxigenin labelled tubulin, and 2 μM of 20% Alexa 647 labelled tubulin in 1 mM GMPCPP, 1mM DTT and BRB80. Reactions were incubated at 37 °C for 1 hour before spun down at 80,000 rpm using a TLA-100 rotor. The Pellet was rinsed twice using warm BRB80 plus DTT and the microtubule seeds were checked using a fluorescent TIRF microscope. The GMPCPP microtubule seeds can be stored at room temperature protected from light for 2 weeks. Labelled tubulin was made by chemical crosslinking of Digoxigenin-NHS ester (Enzo, ENZ-45022) or Alexa647-NHS-ester (Thermo) to microtubules and cycling tubulin using standard protocols, and the final labelling ratio was determined using NanoDrop (Thermo). Tubulin was purified from porcine brains. Tubulin purification and modification was done using standard procedures ^59,60^.

### Dual-trap optical tweezers experiment

Two bead experiments were performed on LUMICKS C-Trap system, which has a microfluidic flow-cell (LUMICKS) containing five laminar flow channels. The system was equipped with a three-colour confocal imaging and a dual optical trap. We used four of the five channels in the C-Trap to perform our experiment. Before the experiment, the flow cell surface was washed with PBS and passivated with 5% Pluronic F-127 for 10 minutes before washing again with Room-temperature BRB80 buffer. Anti-digoxigenin coated polystyrene beads (2 μm diameter, Bangs Lab) were flown into channel 1 and GMPCPP stabilized microtubule seeds labelled with digoxigenin and Alexa647 were flow into channel 2. Channel 3 contained only BRB80 buffer and channel 4 was filled with reaction buffer supplemented with 14 μM unlabelled tubulin, 2 μM of 20% Alexa 647 labelled tubulin, 10 nM CAMSAP3, 100 nM EB3 and 20 nM HSET in 80 mM PIPES, 75 mM KoAC, 15 mM KCl, 1 mM MgCl_2_, 1mM EGTA, 1 mM DTT, 1 mM ATP and 1 mM GTP. During the experiment, the beads were trapped in channel 1 with the dual-optical tweezers at a constant flow generated by ∼0.03 bar pressure, then moved immediately to channel 2 and left to incubate for a few seconds to bind the seeds. Time and concentration of GMPCPP seeds were chosen such that few seeds were attached for the beads as shown in Video S1. Higher concentration of seeds resulted in many more microtubules nucleated at the beads, but were generally not used here (Video S9). The beads were then moved to channel 3 to remove unbound seeds and the seeds coverage was visualized by fluorescent confocal scanning. The beads with microtubule seeds bound were then moved slowly to channel 4. The flow stretched the microtubules growing from the seeds, which allowed to estimate their length. Next, the flow was switched off and the beads were placed at a distance at which microtubules growing form the opposite beads were expected to engage (typically 6 to 12 microns). Force recording was started simultaneously with imaging in three fluorescent channels. Forces were recorded for 10 minutes before flow was switched back on and beads were discarded to start a new round of experiment.

Images were processed Fiji (ImageJ v 2.14) and data was analysed in OriginPro 2021. Direction of force was determined by the increase or decrease in the distance between beads. The rate of microtubule sliding was determined by a linear curve fit of the slope of each force peak.

### Coverslips preparation for the *in vitro* experiments

24✕60 mm glass coverslips (VWR) were cleaned by sonication for 15 minutes each sequentially in Milli-Q water, 100% Acetone, 100% ethanol, 1 M KOH. After cleaning, the coverslips were dried with filtered Nitrogen gas and the glass surface were activated by a plasma cleaner (Corning) for 3 minutes in air at maximum power (3W). The coverslips were then transferred to a clean glass container and submerged in a mixture of 1:20 Dichloromethylsilane:Heptane, then incubated at room temperature for 1 hour. After incubation, the solution was decanted and the hydrophobic coverslips were sonicated for 5 minutes subsequently in chloroform, Milli-Q water, then chloroform again. The Hydrophobic coverslips are air-dried and stored in room temperature.

### Microtubule buckling experiments

Experiments were done using a flow cell assembled by separating silanized hydrophobic coverslips and cleaned glass slides (VWR) by double sided tape (3M). The channels were first incubated with anti-digoxigenin nanobodies (AntiDig-Fab, Roche) followed by passivation using 5% Pluronic F-127 for 15 min and blocked further with 1% BSA in BRB80. 1: 50 dilution of Digoxigenin labelled GMPCPP seeds were pipetted into one end of the flow cell channel and blotted with filter paper on the other end of the channel, the seeds were left incubating in the channel for 1 min before washing two times with 100 μL warm BRB80 buffer. 50 μl of pre-warmed reaction mixture containing 14 μM of unlabelled tubulin, 2 μM of 20% Alexa 647 labelled tubulin, 100 nM EB3 and various concentrations of HSET in 80 mM PIPES, 75 mM KoAC, 15 mM KCl, 1 mM MgCl_2_, 1mM EGTA, 1 mM DTT, 1 mM ATP and 1 mM GTP were flown into the channel immediately before imagining. The flow cells were then mounted onto the microscope stage where the objective was warmed to 37 °C with an objective heater (TOKAI HIT). Images were recorded for 300 seconds at a frame rate of 1 Hz on. Fluorescence was split using OptoSplit (Cairn Research) on two Sona sCMOS cameras (Andor). Three colour channels were recorded simultaneously on two cameras by synchronizing laser shutters with camera’s fire tiggers to produce alternating images of one or the other colour on one of the cameras. Without CAMSAP3, antiparallel microtubule bundles were identified by the direction of HSET movement towards the minus end of each microtubule. Images were later processed and analysed using Fiji (ImageJ v2.14) and MATLAB 2019.

### Analysis of the microtubule shapes and forces

Microtubules were digitized using Fiji (ImageJ v2.14) multipoint function. Minimum seven points were used per single microtubule, and individual frames were analysed independently with 5 frame (5 s) interval between them. Coordinates of the points were then transferred to MATLAB. The shape of the microtubule was interpolated using smoothing spline and the curvature as a function of the position along the microtubule was calculated using:

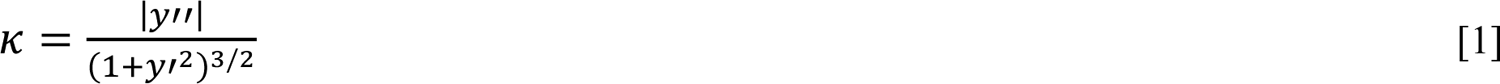

Where *y*(*x*) describes the spline interpolated shape of the microtubule. Microtubule buckling force was calculated as:

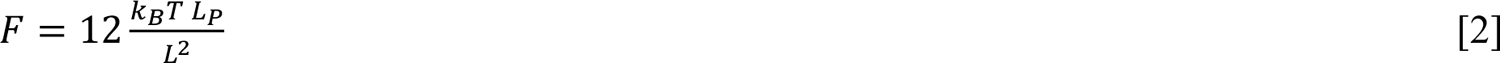

Where *L* is the microtubule contour length and *L*_P_ is the persistence length (we used *L*_P_=4.8mm ^61^). *L* was determined as the length of the microtubule from the tip to the seed at the moment when microtubule became visibly bent. Images of microtubules were digitized and their curvature calculated by custom software written in MATLAB 2019.

### Single-Molecule Fluorescence TIRF microscopy of EB-DNA scaffold movement

Flow cells used in this experiment were assembled and passivated as described in the previous section on microtubule buckling experiment. 50 μl of pre-warmed reaction mixture containing 14 μM of unlabelled tubulin, 2 μM of 20% Alexa 647 labelled tubulin, 1 nM of DNA scaffold EB3 mixture in 80 mM PIPES, 75 mM KoAC, 15 mM KCl, 1 mM MgCl_2_, 1mM EGTA, 1 mM DTT and 1 mM GTP were flow into the channel immediately before imagining. The flow cells were then mounted onto the microscope stage where the objective was warmed to 37 °C. Images were recorded for 300 seconds, at a frame rate of 1 Hz on two Sona sCMOS cameras for dual colour imagining, with an exposure time of 100 ms. Images were later processed and analysed by Image J.

### DNA scaffold force measurement

Since beads were used in these experiments, the flow had to be more tightly controlled. To achieve this, we used flow cells assembled using parafilm sandwiched between a silanized cover slip and a glass slide. The glass slide had metal tubing (New England Small Tube Corp) glued to it on each end using epoxy glue (POXIPOL). Metal connectors were connected by tubing to the syringe pump (Harvard Apparatus). ∼1 nM of biotinylated DNA scaffold coupled to EB3 ensembles were incubated with 1:100 dilution of (1 μm diameter, Bangs Lab) Streptavidin polystyrene beads for 15 min on ice in PBS, pH 7.4 and 2% BSA at a total reaction volume of 50 μl. Beads were then washed twice with ice cold PBS and resuspended in 50 μL PBS. 1 μl of EB3 DNA scaffold ensemble coupled beads were mixed with 50 μl of room-temperature reaction mixture containing 14 μM of unlabelled tubulin, 2 μM of 20% Alexa 647 labelled tubulin, in 80 mM PIPES, 75 mM KoAC, 15 mM KCl, 1 mM MgCl_2_, 1mM EGTA, 1 mM DTT and 1 mM GTP before flowing into the flow cell.

Force measurement was performed by trapping a bead using a single-optical trap (JPK, Nanotracker) and holding it ∼100 nm above the coverslip surface. The bead was moved and placed in front of a growing microtubule and the force were recorded at 2kHz with a typical trap stiffness of 0.02 pN/nm.

To ensure that movement of the bead was due to a single scaffold only, we diluted the scaffolds such that only ∼ 20% of beads interacted with growing microtubule tips. Assuming binding of the scaffolds to beads is a random process governed by Poisson distribution and considering that ∼ 80% of beads did not have a single active scaffold, this suggests that ∼ 18% of all beads in our experiments must have had a single active scaffold and only ∼ 2% have had two of more scaffolds. Thus, out of all signals detected ∼90% must have been generated by the movement of the single scaffold.

Fluorescent imaging at 647 nm wavelength was performed simultaneously with the force recording using a Andor iXon EMCCD camera at 1Hz, 100 ms exposure time. Force curves were later processed using MATLAB 2019 or OriginPro 2021 software. The force curves were first smoothed down to 20 Hz then fitted with sigmoidal logistic function:

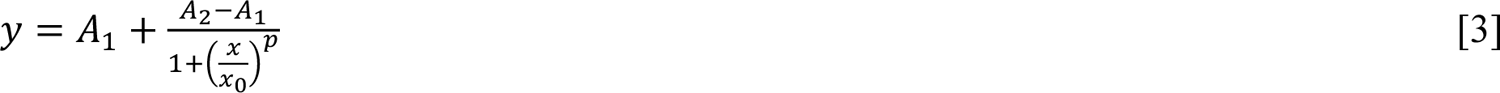

The force was determined as *F* = *A*_1_ – *A*_2_. Only signals in which increase in the force signal agreed with the microtubule growth rate determined from the corresponding fluorescent imaging recording were selected for the further processing.

### Cytosim simulations

The modelling software Cytosim ^62^ was used to model the effect of HSET and EB/HSET system on the organization of two microtubule asters into bipolar structure. Cytosim models microtubules as fibers with rigidity 20 pN·µm/rad and diffusing cross-linking units as particles undergoing Brownian motion in a viscous medium described by over-damped Langevin equations. Viscosity was set in all simulations to 0.01 pN·s/µm^2^. Simulations were run with a time-step of 5 ms for 1000 s. The simulation space was a circle with diameter 60 µm. To mimic the spindle, approximately 100 growing microtubules (0.75 µm in length) were initialized from two asters (diameter 1 µm) separated by 2 µm. Microtubule dynamic instability was simulated as a two-state model defined by a catastrophe frequency (k_cat_ = 0.005 s^-1^ or 0.06 s^-1^ for slower and faster growing microtubules respectively), a rescue frequency (k_rescue_ = 0.06 s^-1^ in all simulations), a constant shrinkage speed (v_s_), and a force-dependent growth speed (v_g_). The growth rate was assumed to be reduced by an antagonistic force (f_a_) by an exponential factor exp (f_a_ / f_g_), where f_g_ is a characteristic growth force which was set to be 1.67 pN. Once a shrinking microtubule’s length is reduced below 100 nm, a new growing microtubule is initialized on that aster.

HSET and EB/HSET units were modelled as two “hands” linked by a Hookean spring. Hands can bind separate microtubules, cross-link the microtubules and apply forces to the microtubules. The association rate was assumed to be k_on_ = 10 s^-1^ for all motors. The dissociation rate was force dependent as: k = k_off_ exp (f_load_ / f_stall_). We used k_off_= 0.1 s^-1^ and assumed f_stall_ either 5 pN or 0.1 pN as discussed in the text. HSET was modelled as a processive motor directed towards the minus end of microtubules. The motor was characterized by its unloaded speed (v_m_) and a stall force (f_stall_). Load forces (f_load_) reduce the motor speed; v = v_m_ (1 + f_load_ ⋅ d / f_stall_). EB was modelled as a plus-end tracking unit, binding specifically to a region near the plus-end of microtubules and tracking the plus-end. Movie frames were generated using the *play* function of Cytosim and assembled into an AVI file in ImageJ. Aster positions were extracted using the *report* function of Cytosim and analysed using a Jupyter notebook. Inter-aster distances were plotted against time.

### Optogenetic inhibition of EB1/HSET interaction on π-EB1 cell lines

π-EB1 H1299 cell lines were a kind gift from Torsten Wittmann. Cells were seeded in a 35mm high glass bottom dish (ibidi) at 80% confluency. The cells were double transfected with pLVX-EB1ΔC-mApple-FRB and pLVX-FKBP-mGFP-HSETΔN using Lipofectamine 3000 (Thermo). Cells were grown in RPMI-1640 media (Gibco) supplemented with 10% FCS (Gibco) and 1% Pen-Strep in a tissue culture incubator supplied with 5% CO2 at 37°C for 24 hours. 1 hour before imaging, the media is replaced with Hanks’ Balanced Salt Solution (HBSS, Gibco) with 10% FCS. 10 µM of MG132 and 0.1% DMSO (Sigma-Aldrich) were added to the media 10 min before imaging for cell cycle arrest at metaphase. For cells treated with Rapamycin, Rapamycin was added to a final concentration of 100 nM in Hanks’ Balanced Salt Solution with 10% FCS and 10 µM of MG132, and the media is added to the cells 10 min before imaging. Imaging was performed no more than 4 hours after MG132 treatment.

Imaging was carried on a Nikon CSU-W1 Spinning Disk confocal system using a 60x 1.49 NA oil immersion lens (Nikon). Cells at metaphase expressing both EB1ΔC-mApple-FRB and FKBP-mGFP-HSETΔN were selected by equal intensity of both red and green fluorescence. For blue light illumination, time-lapsed imaging acquisition was carried out at 488 nm and 561 nm wavelength, with a 0.1 second exposure time. Blue-light illumination was triggered by camera with a 1 second delay after camera exposure and 0.4 seconds blue light intervals, z-stack were taken in the range of 15 µm and 1 µm per slice.

Images were further processed in Image J. The length of the spindle was determined from the pole-to-pole distance using all optical sections. The first timepoint of each time-lapsed video was taken before blue-light stimulation (in the dark) and the rest of the sequence was taken in the presence of the blue light.

## Acknowledgments

We thank all our laboratory members for discussions and critical comments on the manuscript. We thank Thomas Surrey for supporting initial stages of this project, for providing mGFP-EB3 and mCherry HSET plasmids and protocols for tubulin purification from porcine brain and the helpful discussions and advice. We thank Torsten Wittmann for sharing π-EB1 cell lines and for helpful discussions and assistance in the project. We thank Anna Akhmanova for kindly providing plasmids for CAMSAP3. We thank Rocco D’Antuono, Matt Renshaw, Gregory Mashanov and the rest of the Crick Science Technology Platforms for their assistance in this project. This work was supported by the Francis Crick Institute, which received funding from the UK Medical Research Council (FC001750), Cancer Research UK (FC001750) and the Wellcome Trust (FC001750) through “Mechanobiology and biophysics” award to MM.

## Author Contributions

LC designed, performed and analysed experiments; GP assisted with force spectroscopy measurements; DS performed Cytosim simulations; JG supported initial stages of the project and established of biochemical procedures. SC co-supervised DS. LC and MM wrote the manuscript. MM designed and supervised the research.

## Open Access

For the purpose of Open Access, the authors have applied a CC BY public copyright licence to any Author Accepted Manuscript version arising from this submission.

## Supplementary Figure Legends

**Figure S1.**
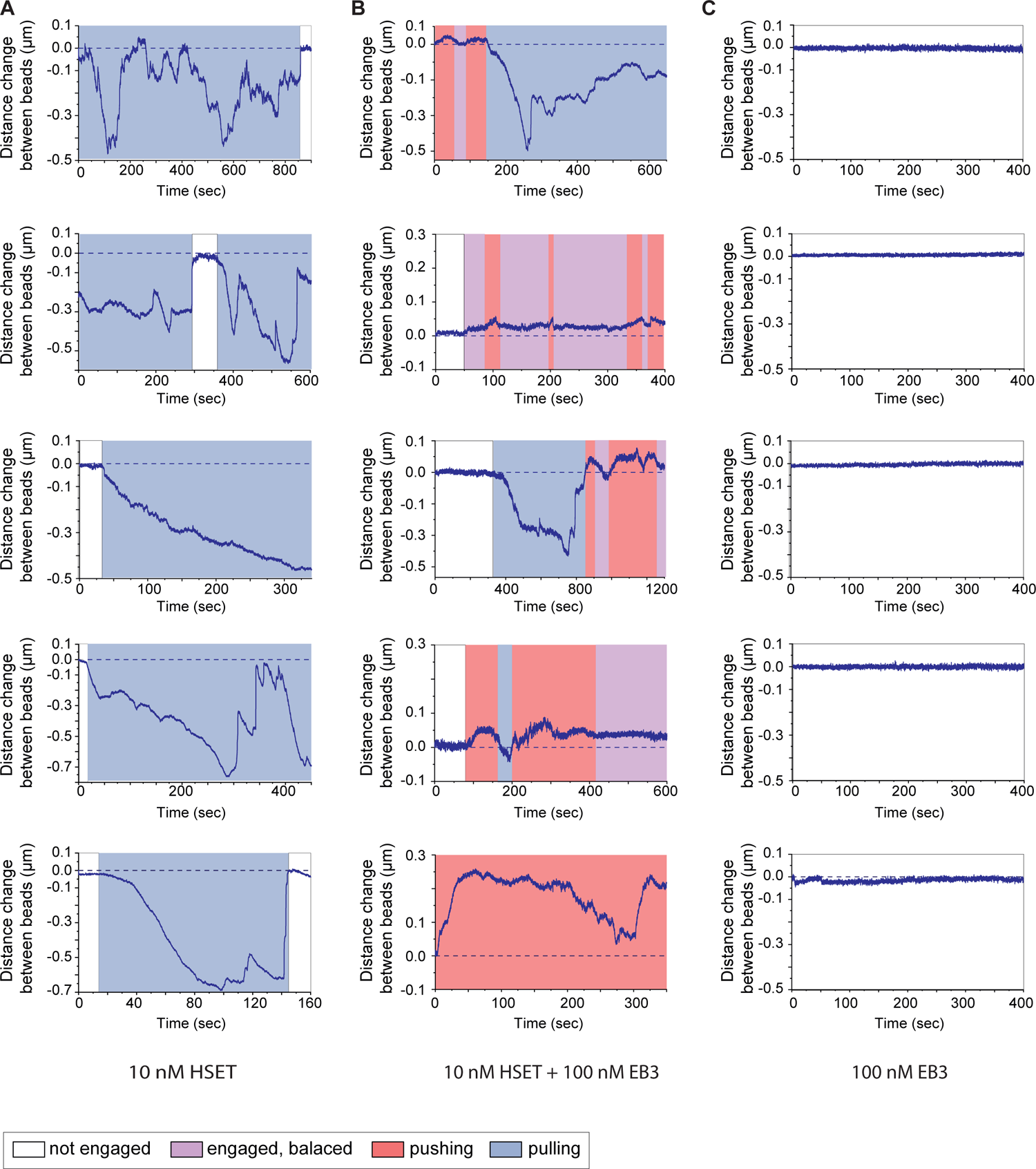
Additional examples of traces for the double bead “artificial spindle” assay. **A**, Additional examples of traces in the presence of HSET alone. **B,** Additional examples of traces in the presence of HSET and EB. Colour shows where the system is experiencing pushing and pulling forces and where it is balanced. **C,** Examples of traces in the presence of EB alone. No force detected and asters do not visibly engage. In A-C only the distance between the optically trapped beads is shown (force is linearly proportional to the distance).

**Figure S2.**
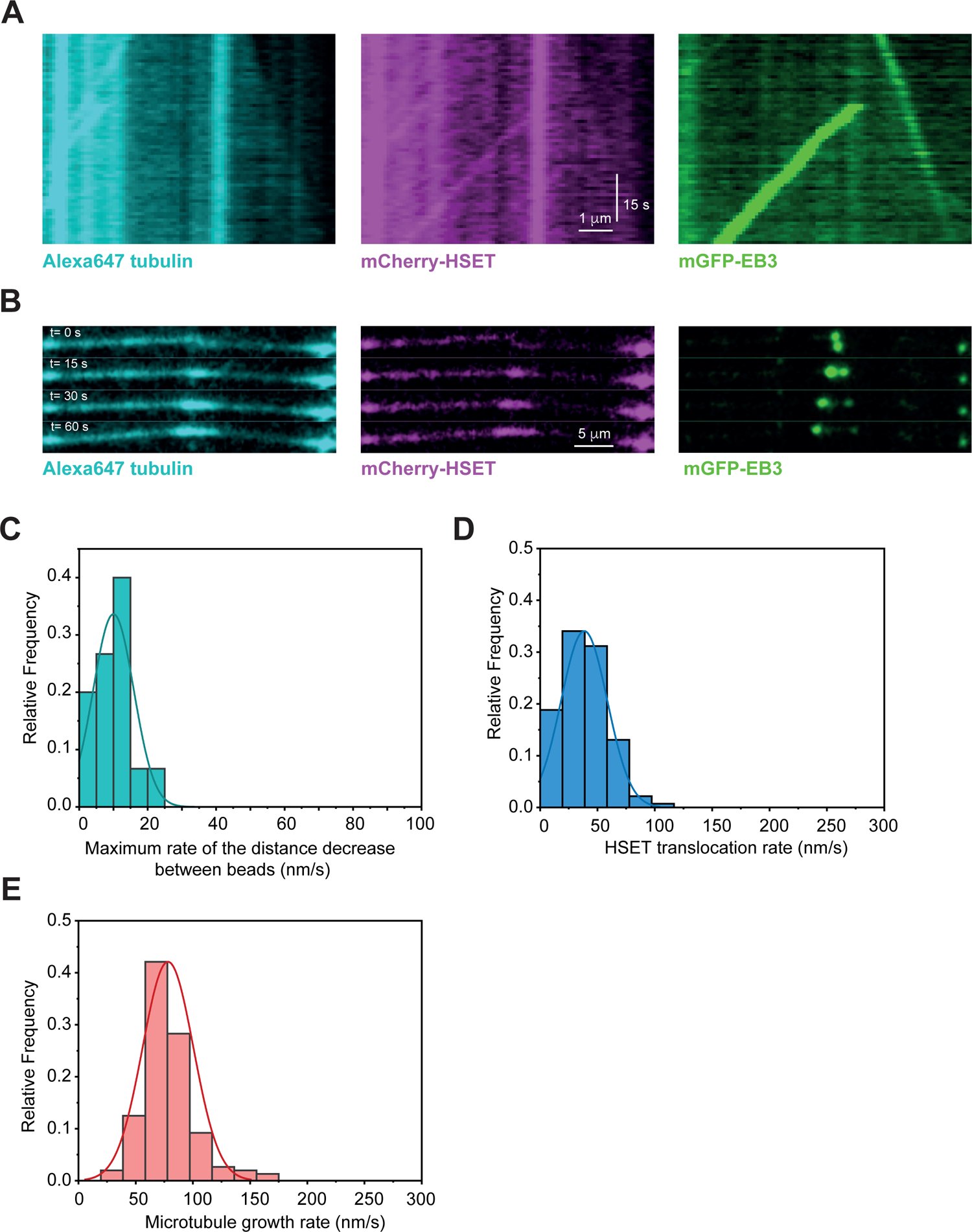
Quantification of HSET and microtubule dynamics. **A**, Kymograph shows translocation of EB molecules by HSET towards microtubule minus ends indicating the interaction between EB and HSET. **B**, Still images of individual channels are shown for the buckling microtubules from example in Figure 2A. **C,** Maximum rate of the decrease between the two optically trapped bead driven by HSET action in the antiparallel microtubule overlaps (no EB present). **D,** Speed of individual HSET translocation towards microtubule minus ends along individual microtubules measured in TIRF assay. **E,** Individual microtubule growth rate in the buckling experiments.

**Figure S3.**
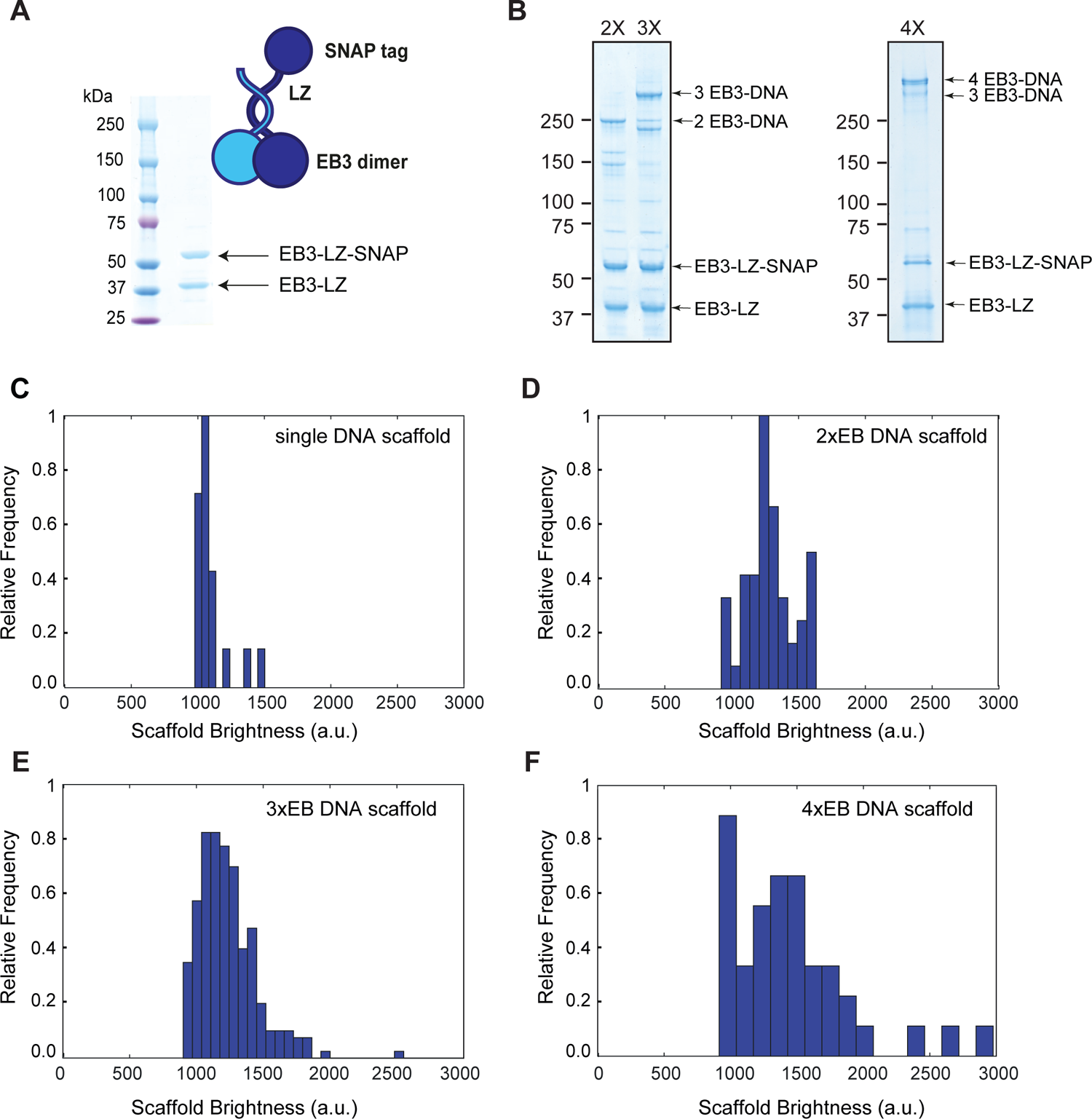
Characterization of the DNA scaffolds. **A**, Denaturing SDS-PAGE of the purified EB3-LZ-SNAP/EB3-LZ heterodimers. The heterodimer with a single SNAP tag ensures that it can attach only to a single position of DNA backbone. **B,** SDS-PAGE (not boiled/heated) of the EB3 ensembles coupled to DNA. 2x, 3x and 4x indicate samples with 2,3 and 4 EBs per scaffold correspondingly. SDS in the gel separates EB3 heterodimers. The sample was not heated, which prevented melting of the dsDNA structure and kept the scaffold intact. The major top bands in each case (arrows) correspond to scaffolds with the expected number of attached EBs. **C,** Control experiment in which individual DNA scaffolds were adsorbed directly on the surface of the coverslip and brightness of the individual Cy3 dyes associated with single scaffolds were quantified. **D,** Brightness of 2xEB DNA scaffolds that were used to determine the run lengths of the 2xEB complexes on growing microtubule tips. Their brightness corresponds to the single Cy3 fluorophore from A. **E – F,** Same as in B, but for scaffolds with 3 or 4 EB dimers respectively.

**Figure S4.**
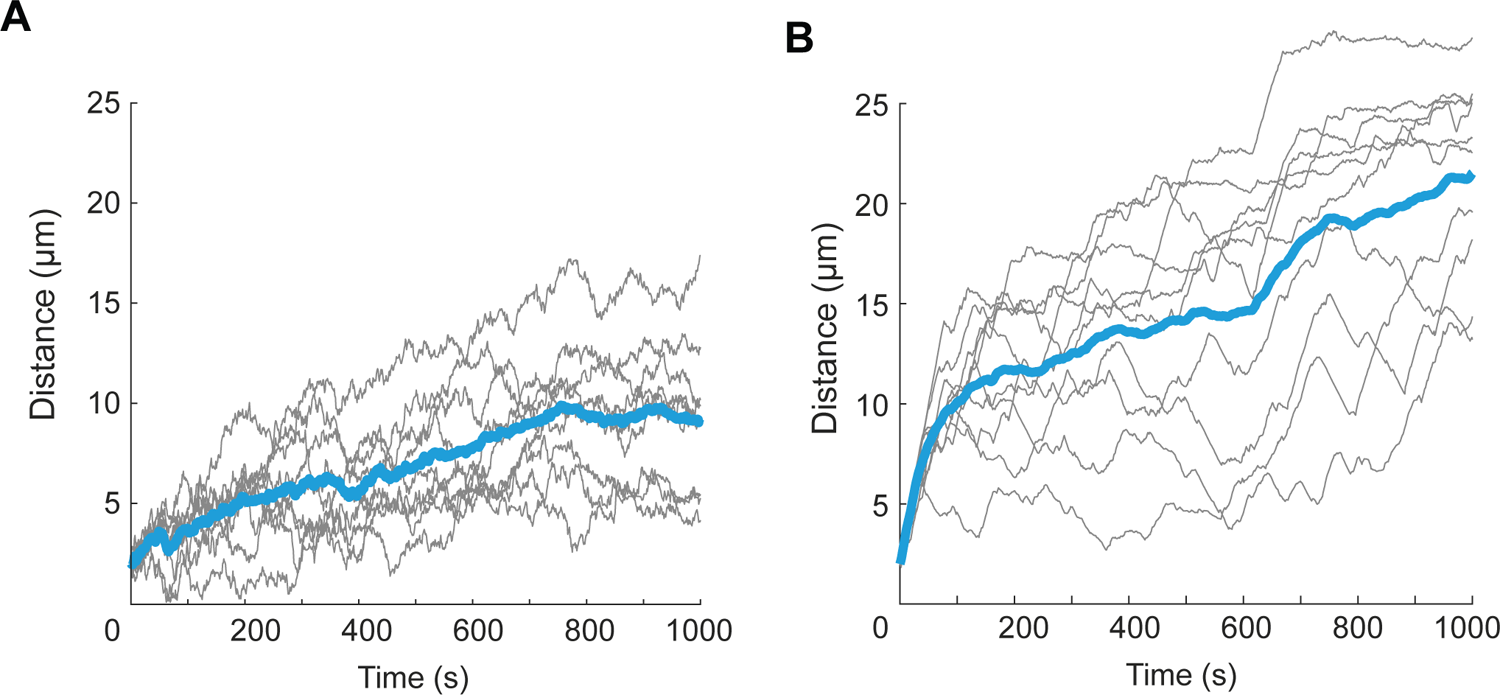
Additional simulations of the pole-to-pole distance between two asters of dynamic microtubules interacting via HSET and EB/HSET. **A**, Simulation in which total number of HSET and EB/HSET complexes was reduced from 1200 to 60. Individual traces experience more noise from increased stochasticity but the steady state distance between poles remains the same as in Figure 4G indicating that it does not depend on the absolute number of motors. **B,** Simulation in which speed of microtubule growth was 4x higher than the speed of the HSET movement (250 nm/s and 60 nm/s respectively). Simulations show increased distance between poles due to longer microtubules and their faster growth rate.

**Figure S5.**
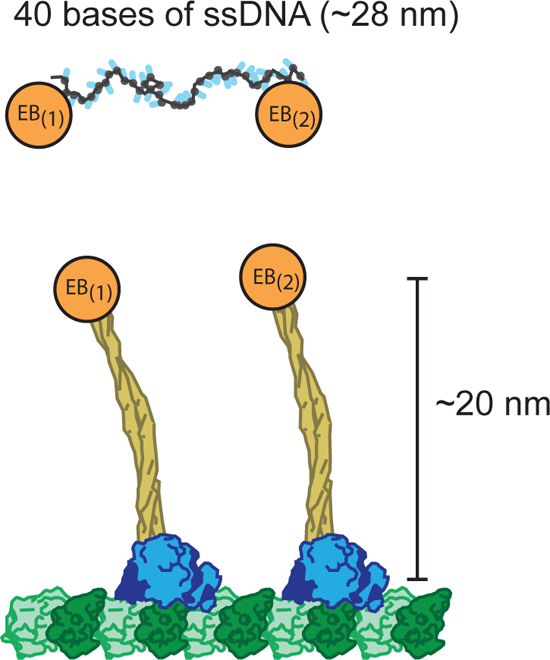
Geometrical comparison of the EB binding between DNA scaffolds and microtubule/HSET. Schematics of two EB molecules coupled to a flexible ssDNA scaffold. Distance between two neighbouring EB binding sites is ∼25 nm in contour length. Two HSET molecules bound to microtubule lattice are shown in scale with DNA. The coiled-coil region of the HSET is ∼ 20 nm. Distance between two HSET as shown is 16 nm (two tubulin dimers), but can be different depending on the HSET positions.

## Supplementary Video Legends

**Video S1.** Dual trap optical tweezers experiment with 10 nM HSET. Video showing crosslinking of antiparallel microtubule by HSET in double-bead optical trap with low density of GMPCPP seeds. Alexa647 microtubule seeds are coloured by magenta, and mCherry-HSET is coloured by green, the two trapped beads are coloured by blue.

**Video S2.** HSET transports EB to the microtubule minus ends. HSET interaction with EB3 allows EB3 to be transported to the microtubule minus end by HSET. For corresponding kymograph see Supplementary Figure S2.

**Video S3.** Two antiparallel microtubules buckle in the presence of EB3 and HSET. related to Figure 2A.

**Video S4.** Simulation of the two aster dynamics driven by HSET. In this simulation fraction of EB/HSET was 10% and dynamics was dominated by the action of HSET alone that led to the two asters fusion. Colour legend: White – microtubules, Magenta – unbound HSET, Dark blue – unbound EB/HSET, Pink – bound HSET, Light blue – bound EB/HSET.

**Video S5.** Simulation of the two aster dynamics driven by EB/HSET. In this simulation fraction of EB/HSET was 90%. In these case asters stabilize in bipolar configuration. Colours follow Video S4.

**Video S6.** Dissociation of π-EB1 from the microtubule plus-ends by blue-light stimulation in π-EB1 H1299 cell lines. Control experiment shows disappearance of EB1 comets after blue-light activation in interphase cells.

**Video S7.** Active spindle shortening in π-EB1 H1299 cell lines without Rapamycin treatment after blue-light activation. EB1ΔC-mApple-FRB is in red, and FKBP-mGFP-HSETΔN is in green. Related to Figure 5B.

**Video S8.** Mitotic π-EB1 H1299 cell after rapamycin treatment and blue-light activation. EB1ΔC-mApple-FRB is in red, and FKBP-mGFP-HSETΔN is in green. Related to Figure 5B.

**Video S9.** Example of a dual trap optical tweezers experiment with 10 nM HSET with higher density of microtubule seeds than in Video S1. **Notations follow Video S1.**

